# Protective heterologous T cell immunity in COVID-19 induced by MMR and Tdap vaccine antigens

**DOI:** 10.1101/2021.05.03.441323

**Authors:** Vijayashree Mysore, Xavier Cullere, Matthew L. Settles, Xinge Ji, Michael W. Kattan, Michaël Desjardins, Blythe Durbin-Johnson, Tal Gilboa, Lindsey R. Baden, David R. Walt, Andrew Lichtman, Lara Jehi, Tanya N. Mayadas

**Affiliations:** Department of Pathology, Brigham and Women’s Hospital & Harvard Medical School, Boston, MA, 02115, USA; Bioinformatics Core Facility in the Genome Center, University of California Davis, Davis, CA, 95616, USA; Quantitative Health Science Department, Cleveland Clinic, Cleveland, OH 44195, USA; Department of Medicine, Brigham and Women’s Hospital & Harvard Medical School, Boston, MA, 02115, USA; Division of Biostatistics, University of California Davis, Davis, CA 95616, USA; Neurological Institute, Cleveland Clinic, Cleveland, OH 44195, USA

**Keywords:** SARS-CoV-2, heterologous immunity, antigens, vaccines, memory T cells, TCR repertoire, transcriptome, antigen presenting cells, risk prediction

## Abstract

T cells are critical for control of viral infection and effective vaccination. We investigated whether prior Measles-Mumps-Rubella (MMR) or Tetanus-Diphtheria-pertussis (Tdap) vaccination elicit cross-reactive T cells that mitigate COVID-19. Using co-cultures of antigen presenting cells (APC) loaded with antigens and autologous T cells, we found a high correlation between responses to SARS-CoV-2 (Spike-S1 and Nucleocapsid) and MMR and Tdap vaccine proteins in both SARS-CoV-2 infected individuals and individuals immunized with mRNA-based SARS-CoV-2 vaccines. The overlapping T cell population contained effector memory T cells (TEMRA) previously implicated in anti-viral immunity and their activation required APC-derived IL-15. TCR- and scRNA-sequencing detected cross-reactive clones with TEMRA features among the cells recognizing SARS-CoV-2, MMR and Tdap epitopes. A propensity-weighted analysis of 73,582 COVID-19 patients revealed that severe disease outcomes (hospitalization and transfer to intensive care unit or death) were reduced in MMR or Tdap vaccinated individuals by 38-32% and 23-20% respectively. In summary, SARS-CoV-2 re-activates memory T cells generated by Tdap and MMR vaccines, which may reduce disease severity.

## INTRODUCTION

T cells are essential for early control of acute viral infection and for generation of B cells producing protective antibodies. CD4+ helper T cells (Th) induce B cells to produce high affinity antibodies to viral protein antigens. Effector CD8+ T cells (cytotoxic T lymphocytes, CTL) and CD4+ T cells eradicate infected, virus-producing cells via direct killing or secretion of cytokines such as interferon-γ (IFN-γ), which enhances inflammatory functions that support viral clearance. Antigen presenting cells (APC) such as classical dendritic cells (DC) play a critical role in initiating the cellular immune response by processing and presenting internalized antigen to T cells, which become activated and proliferate. T cell expansion following productive immunity usually produces a memory T cell population that can persist for decades. Compared to their naïve precursors, memory T cells are more abundant, have a lower threshold for activation and rapidly reactivate effector functions following antigen encounter. They are also maintained in barrier tissues to rapidly respond to reinfection. Thus, a major goal of vaccines is the induction of strong and durable T and B cell memory (*1*).

The appearance of SARS-Co-V-2 specific CD4+ and CD8+ T cells early after symptom onset (*2-5*) is associated with rapid viral clearance and mild disease (*3*), whereas delayed T cell responses is correlated with worse clinical outcomes (*6*). Antigen-specific T-cell responses evaluated by exposing peripheral blood mononuclear cells (PBMC) to peptide pools (*7-10*) suggest that Spike (the target of most COVID-19 vaccines), nucleocapsid and M envelope proteins are the most relevant CD4+ and CD8+ T cell targets (*7, 8, 10-12*). On the other hand, peptide-MHC tetramer staining, used to screen epitopes for T cell recognition across different HLA alleles, found that CD8+ T cells specific to nucleocapsid were present at a higher frequency than those specific for Spike-or non-structural proteins (*13, 14*). In several studies, the magnitude of SARS-CoV-2-specific IgG and IgA titers correlated with the SARS-CoV-2 T cell response (*10, 11, 15*). Interestingly, memory T cells specific for related coronaviruses including those that cause the common cold that cross-react with SARS-Cov-2 antigens have been detected in a large percent of unexposed individuals (*7, 8, 11, 16, 17*). Moreover, profiling of the TCR repertoire of T cells isolated from unexposed and SARS-CoV-2 convalescent patients and expanded *in vitro* with predicted immunodominant viral peptides show clonal expansion of T cells with TCR sequences recognizing peptides from other viruses including HCMV, HHV-5 and influenza A (*18*). The impact of these pre-existing, cross-reactive memory T cells on disease outcomes are largely unknown (*19*).

A diverse T cell response is essential to provide early control of acute viral infection. Thus, a major goal of vaccines is the induction of strong and durable B and T cell memory (*1*). Adaptive heterologous immunity is a response to a microbe mediated by memory T cells generated against the antigens of a different microbe, either through infection or immunization (*20*) and may provide enhanced immunity to novel pathogens. The foundation of adaptive heterologous immunity during infection is the existence of cross-reactive T or B cell clones to epitopes present in two different pathogens. Yet, to our knowledge, definitive identification of the relevant epitopes or the cross-reacting lymphocyte clones in humans has not been previously achieved. Here, we leveraged a sensitive new assay for antigen-specific T cell responses to physiologically processed antigens to determine if SARS-CoV-2 protein-specific T cells in the blood of convalescent SARS-CoV-2 infected or uninfected SARS-CoV-2 mRNA vaccinated individuals are cross-reactive with the antigens in MMR (Measles-Mumps-Rubella) and Tdap (Tetanus-Diphtheria-Pertussis) vaccines, known to be highly effective in eliciting long-lasting protective immunity (*21*) and therefore T and B cell memory responses. Next, we used a large, deeply phenotyped COVID-19 cohort adjusted for multiple patient characteristics to determine if prior trivalent MMR or Tdap vaccinations attenuates disease severity in COVID-19 patients.

## RESULTS

### Correlation of T cell responses to SARS-CoV-2 and Tdap and MMR vaccine antigens in COVID-19 infected subjects

The strategy used to assess the correlation of T cell responses to SARS-CoV-2 and Tdap and MMR vaccine antigens is summarized in **Fig. 1A**. Subjects with PCR-confirmed SARS-CoV-2 infection and uninfected controls (confirmed by absence of SARS-CoV-2 antibodies) were included in the study (**Table 1**). Plasma cytokine profiles were similar in both groups (fig. S1A), ruling out their potential contribution to observed results. Prior studies of human T cell responses *in vitro* have primarily relied on DCs derived from monocytes (*22, 23*) or PBMCs (*24*) that contain a mixture of poorly immunogenic APCs. Here, we generated highly immunogenic APCs expressing CD11c, HLA-DR, T cell co-stimulatory molecules and the migration receptor CCR7 (fig. S1B) by engaging the FcγRIIIB receptor on neutrophils with a complexed anti-FcγRIIIB antibody, referred to as AAC (see methods) and culturing them in GM-CSF (*25*); hereafter, these cells are referred to as neutrophil derived APCs (nAPC). nAPCs were pulsed with native SARS-CoV-2, MMR or Tdap antigens and co-cultured with autologous T cells on interferon-γ (IFN-γ) ELISpot microplates for only 18hrs to limit antigen mediated T cell proliferation and phenotypic changes. IFN-γ is a robust marker of memory T cell activation, whereas it is low to undetectable in TCR activated naive T cells (*26, 27*). AAC-treated neutrophils pulsed with Spike-S1 induced robust T cell activation only in SARS-CoV-2-infected individuals (**Fig. 1B**, fig. S1C). No IFN-γ-secreting T cells were observed when nAPCs were not pulsed with antigen, with isotype antibody treated neutrophils pulsed with antigen, and with T cells incubated with antigen alone (fig. S1C). Most COVID-19 patients seroconvert within 7-14 days of infection (*28, 29*) and IgG antibodies persist for at least 9 months after exposure (*30*). SARS-CoV-2 specific IgG titers detected by high-resolution profiling of plasma (*31*) were present in all patients screened 14 days or more after a positive SARS-CoV-2 test (fig. S1D), whereas antibodies were observed in only a subset of patients within 2-6 days of a SARS-CoV-2 positive test or 5-11 days of developing symptoms (**Fig. 1C**). By contrast, T cell responses were detected at all time points following infection (**Fig. 1B, Table 1**), suggesting that effector memory T cells reactive to SARS-CoV-2 antigens predate COVID-19 infection.

**Figure 1:**
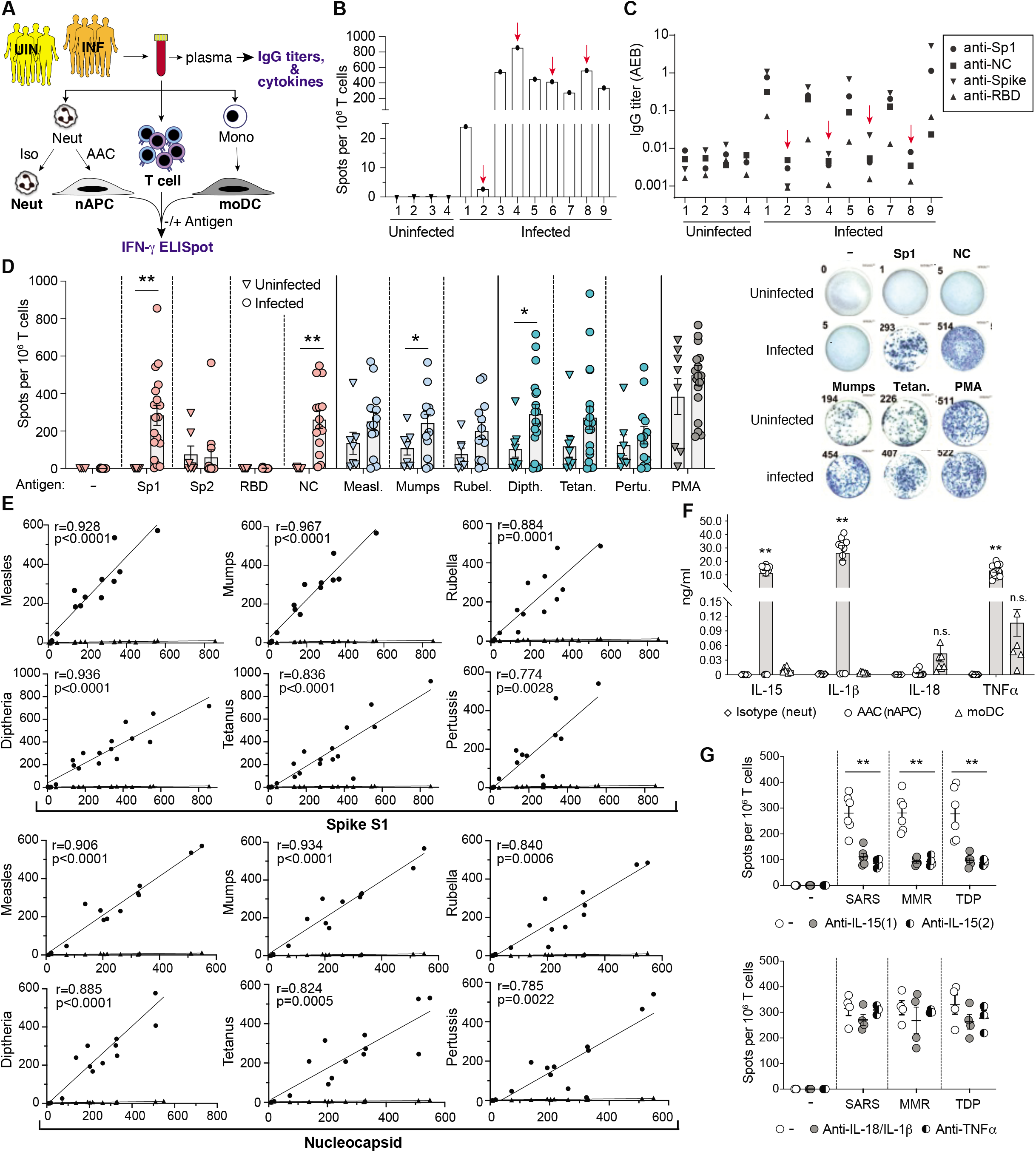
T cell responses to SARS-CoV-2, MMR and Tdap antigens in SARS-CoV-2 infected and uninfected donors. **A)** Blood was collected from uninfected (UNI) and PCR-confirmed, SARS-CoV-2 Infected (INF) donors. Plasma was analyzed for antibodies to SARS-CoV-2 antigens and cytokines. Blood was divided and treated with Isotype control or AAC for 2hrs and neutrophils were then isolated and cultured in GM-CSF for 2 days. AAC treated neutrophils converted to APCs (nAPC) while Isotype treated controls remained neutrophils. PBMC were harvested to isolate CD3^+^ T cells and monocytes, which were cultured in cytokines to generate dendritic cells (moDC). Neutrophils (Isotype), nAPC (AAC) and moDC were loaded with antigens and co-cultured with autologous T cells on IFN-γ ELISpot plates for 18hrs. **B)** nAPC from unexposed (1-4) and exposed (1-9) were co-cultured with autologous CD3 T cells and Spike-S1 on IFNγ-ELISpot plates. The number of spots per 1 × 10^6^ cells is reported. **C)** IgG titers in sera of unexposed and exposed donors to indicated SARS-CoV-2 antigens. Red arrows in B)-C) identify samples with high frequency of IFN-γ^+^ T cells but no detectable SARS-CoV-2 antibodies. **D)** nAPCs generated from 8 unexposed and 18 exposed donors were loaded with indicated individual SARS-CoV-2, MMR and Tdap antigens and analyzed for T cell responses as in A) *(left panel)*. Representative images of wells with IFN-γ^+^ spots from an ELISpot assay are shown (*right panel*). *p<0.05; **p<0.005 by two-tailed Mann-Whitney test with Bonferroni correction for multiple comparisons. **E)** A correlation of Spike-S1 or Nucleocapsid derived IFN-γ^+^ spots with indicated vaccine antigens (circles) and percent of nAPCs generated (diamonds) in exposed donors was conducted using Spearman’s rank correlation coefficients (r). *p<0.05; **p<0.005 using two-way analysis of variance and Bonferroni’s multiple comparison test. **F)** Cytokine levels detected in the supernatants of neutrophils treated with isotype or AAC (nAPC) and monocyte derived dendritic cells (moDC) cultured for 72hrs.**p<0.005 using two-way analysis of variance and Bonferroni’s multiple comparison test. **G)** As in B), ELISpot assays measuring IFN-γ secretion by T cells co-cultured with nAPCs pulsed with combined SARS-CoV-2 (Spike S1, Nucleocapsid, RBD), MMR (Measles, Mumps and Rubella) or Tdap (Tetanus, Diphtheria and Pertussis) antigens were evaluated in the presence of two independent anti-IL15 antibodies (*top panel*), anti-IL-1β/anti-IL18 or anti-TNFα (*bottom panel*). The number of spots per 10^6^ T cells was quantified for each antigen combination. Two-way analysis of variance and Dunnett’s multiple comparison test was used.**p<0.005. All Data expressed are mean ± sem.

**Table 1:** Characteristics of COVID-19 Infected and Uninfected Donors for cellular and molecular assays

| Infected |  |  |  |  | Uninfected |  |  |
| --- | --- | --- | --- | --- | --- | --- | --- |
|  | M/F | Age | Days from first COVID+ test to Blood draw | Days from first Symptoms to Blood draw |  | M/F | Age |
| 1 | M | 74 | 22 | 11 | 1 | F | 32 |
| 2 | F | 89 | 4 | 5 | 2 | F | 27 |
| 3 | M | 52 | 37 | 36 | 3 | M | 38 |
| 4 | F | 61 | 5 | 5 | 4 | F | 35 |
| 5 | F | 25 | 94 | 98 | 5 | F | 58 |
| 6 | F | 25 | 6 | 11 | 6 | M | 26 |
| 7 | M | 36 | 69 | NK | 7 | M | 67 |
| 8 | F | 47 | 100 | 109 | 8 | M | 56 |
| 9 | F | 68 | 2 | 5 |  |  |  |
| 10 | M | 72 | 5 | 9 |  |  |  |
| 11 | F | 52 | 124 | 133 |  |  |  |
| 12 | F | 37 | 94 | 96 |  |  |  |
| 13 | F | 51 | 120 | 120 |  |  |  |
| 14 | F | 64 | 131 | 139 |  |  |  |
| 15 | M | 32 | 204 | NK |  |  |  |
| 16 | M | 54 | 13 | 13 |  |  |  |
| 17 | M | 52 | 96 | NK |  |  |  |
| 18 | F | 50 | 112 | NK |  |  |  |

NK: not known

Uninfected, SARS-CoV-2 vaccinated
| M/F | Age | Days from second dose of vaccine to Blood draw |
| --- | --- | --- |
| F | 32 | 74 |
| M | 67 | 80 |
| F | 35 | 71 |

Donors for V(D)J clonotyping and scRNAseq studies
| Infected |  |  |  | Uninfected |  |
| --- | --- | --- | --- | --- | --- |
| M/F | Age | Days from first COVID+ test to Blood draw | Days from first Symptoms to Blood draw | M/F | Age |
| F | 25 | 192 | 196 | F | 32 |
| M | 52 | 156 | 155 | M | 67 |
| M | 32 | 204 | NK | F | 35 |

To elucidate whether SARS-CoV-2 infection results in the activation of memory T cells generated by prior MMR or Tdap vaccination, we examined T cell responses to a broader range of SARS-CoV-2 antigens and antigens present in the MMR and Tdap vaccines and compared results between infected individuals, the majority of who were convalescent patients (**Table 1**) and uninfected individuals. Robust T cell responses to Spike-S1 and Nucleocapsid were observed in infected individuals. Those to Spike-S2 were low and variable, consistent with prior reports (*16, 17*) while no response to receptor binding domain (RBD) (within Spike-S1) was detected. Responses to MMR and Tdap antigens were detected in all individuals but the frequency of T cell responses to MMR and Tdap antigens trended higher in the infected versus the uninfected group (**Fig. 1D**). Importantly, T cell responses to Spike-S1 and Nucleocapsid and individual MMR or Tdap vaccine antigens were strongly correlated in individuals who were previously infected with SARS-CoV-2 (**Fig. 1E)**. For a subset of blood samples from infected and uninfected subjects, DCs were derived from monocytes (moDC), loaded with SARS-CoV-2 and vaccine antigens and assessed for IFN-γ secreting autologous T cells. The overall T cell response to DCs was markedly lower (fig. S1E) than for nAPCs (**Fig. 1D**) despite comparable expression of HLA-DR and T cell co-stimulatory molecules (fig. S1F). Nonetheless, a significant, albeit weaker correlation of T cell responses to Spike-S1 and nucleocapsid with MMR antigens and Pertussis was observed (fig S1G).

### Superior T cell activation by nAPCs is related to IL-15 production

Higher T cell activation by nAPCs versus moDCs may reflect differences in expression of immunomodulatory cytokines (*32*) that sensitize cross-reactive T cells to non-cognate antigen. For example, IL-15 enhances memory CD8 TCR affinity and avidity (*33-35*), promotes IFN-γ production by CD8 T_EMRA_ (*36*) and counteracts CD4 T cell suppression by T regulatory cells (*37*). nAPCs secreted log fold higher amounts of IL-15, IL-1β, and TNFα compared to moDC; total levels were comparable between uninfected and infected individuals (**Fig. 1F**). To investigate the contribution of these cytokines on T cell activation, we treated nAPC and T cell co-cultures with neutralizing antibodies. Two IL-15 antibodies significantly reduced the number of IFN-γ secreting T_EM_ and TEMRA cells, whereas combined IL-1β plus IL-18 antibodies and a TNFα blocking antibody had no effect (**Fig. 1G**). A similar analysis of moDCs showed that blocking IL-15 led to minimal reduction in T cell activation (fig. S1H).

### Phenotype of Memory T cells reactive to SARS-CoV-2, MMR and Tdap

To characterize the CD4^+^ and CD8^+^ T cell lineages activated by SARS-CoV-2, MMR and Tdap antigens in infected individuals, we used flow cytometry to assess cell surface markers for naïve (TNAIVE), central memory (TCM), effector memory (TEM) and effector memory re-expressing CD45RA (TEMRA) T cells (gating strategy, fig. S2A, B) and markers of activation, homing, function and proliferation in T cells positive for intracellular IFN-γ (*38*). We observed a significant increase in the frequency of TEMRA markers in CD4+ and CD8+ T cells co-cultured with nAPCs loaded with SARS-CoV-2, Tdap or MMR antigens compared to T cells co-cultured with non-antigen loaded nAPCs (**Fig. 2A, B**). The CD4 TEMRA expressed surface markers associated with cytotoxicity, GPR56 (*39*) and migration into infected tissue, CX3CR1 (*40*) (**Fig. 2A**) and the CD8 TEM and TEMRA exhibited an increase in the activation marker CD69 (**Fig. 2B**). A small degree of proliferation (2.87±0.12% of total CD3^+^ T cells) was also present. IL-15 blockade caused a significant decrease in the percentage of IFN-γ^+^ T_EMRA_ among CD4+ T cells and IFN-γ^+^ T_EM_ and T_EMRA_ among CD8+ T cells (**Fig. 2C**). In T cells co-cultured with SARS antigen loaded moDC, IFNγ producing CD4 T_EM_ and a very small population of CD4 or CD8 TEMRA were detected; anti-IL-15 had no effect on the activation of either of these populations (fig. S2C). Notably, T_EMRA_ result from antigen reactivation of effector memory T cells, are implicated in protective anti-viral immunity (*39-43*) and are prevalent in convalescent COVID-19 patients (*44*).

**Figure 2:**
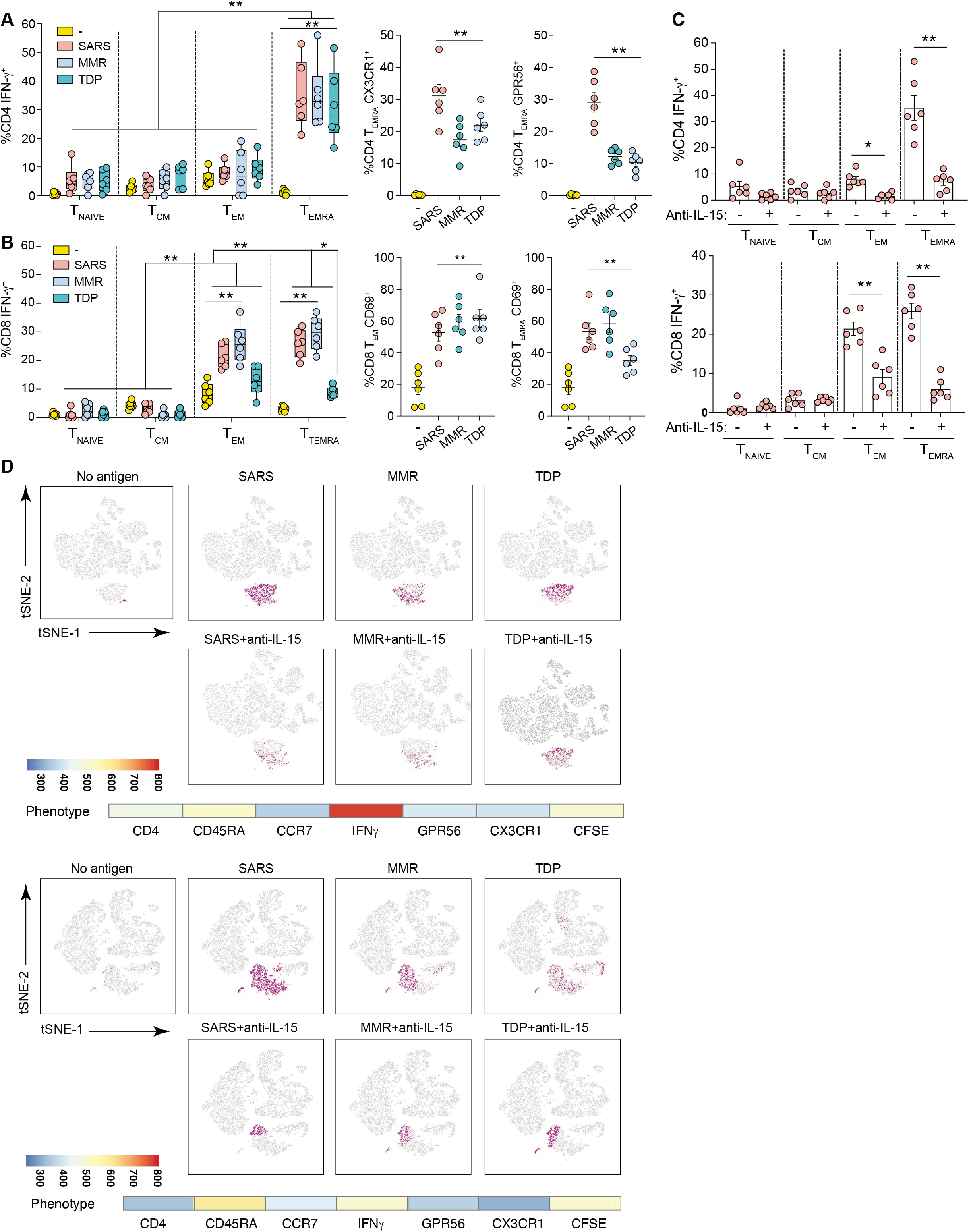
Phenotype of T cells responding to SARS-CoV-2, MMR and Tdap antigens. **A-D)** T cells from the ELISpot wells were harvested after 18h co-culture with nAPCs. The cells were treated with Brefeldin A for 5h, stained for phenotypic surface and intracellular markers and assessed by flow cytometry. **A)** Proportions of naïve (TNAIVE CCR7^+^CD45RA^+^), central memory (TCM CCR7^+^CD45RA^-^), effector memory (TEM CCR7^-^CD45RA^-^) and effector memory RA (TEMRA CCR7^-^CD45RA^+^) CD4 T cells expressing IFN-γ *(left panel)* and additional cytolytic markers, CX3CR1 and GPR56 in TEMRA are shown *(right panel)*. **B)** CD8 T cells expressing IFN-γ were analyzed and classified as TNAIVE (CCR7^+^CD45RA^+^CD27^+^), TCM (CCR7^+^CD45RA^-^CD27^+^), TEM (CCR7^-^CD45RA^-^CD27^-^) and TEMRA (CCR7^-^CD45RA^+^CD27^-^) *(left panel)*. TEM and TEMRA were further evaluated for CD69 (*right panel*). A and B were analyzed using three-way analysis of variance and Tukey’s multiple comparisons test. *p<0.05, **p<0.005. **C)** Anti-IL-15 or vehicle control was included during T cell co-cultures with nAPCs pulsed with SARS-CoV-2 antigens and subsets of CD4 (*top panel*) and CD8 (*bottom panel*) T cells producing IFN-γ were determined. Two-way analysis of variance and Dunnett’s multiple comparisons test was used. *p<0.05, **p<0.005. **D)** FLOWsom based visualization on tSNE plots of flow cytometry datasets for CD4+ live T cells from 2 individual exposed donors. Cells shown in grey correspond to total CD4 T cells from down-sampled and concatenated specimens stimulated with SARS-CoV-2 (Spike S1, Nucleocapsid, RBD), MMR and Tdap antigens with each in the presence or absence of anti-IL-15 antibody, to create a single tSNE map. Cell populations defined by FLOWsom for each antigen were then projected onto tSNE space and a population was identified that overlapped between SARS-CoV-2 (SARS), MMR and Tdap antigens (purple). The phenotype of the overlapping population between the antigens was defined by evaluating the following markers, CD4, CD45RA, CCR7, CD27, GPR56, CX3CR1, IFN-γ and shown as heatmaps along with a continuous scale.

To further define the T cell populations reactive to nAPCs loaded with the three antigens, we visualized the profile of CD4+ T cells using viSNE, which uses t-stochastic neighbor embedding (t-SNE) to generate a two dimensional map of cell relatedness based on marker profile similarity (*45*). viSNE depicted an IFN-γ^+^ T cell cluster that was responsive to SARS-CoV-2, MMR or Tdap antigens and had features of TEMRA (**Fig. 2D**, fig. S2D). The frequency of CD4+ TEMRA cells ranged from 5 to 13%, which is consistent with the reported variability in the frequency of TEMRA (<0.3-18% of total CD4 T cells) even in the absence of infection (*46*). The cluster containing CD4+ TEMRA was significantly reduced after anti-IL15 treatment (**Fig. 2D**). An overlapping region among the three sets of antigen was not dectected for CD8+ T cells (not shown), although inclusion of antibodies to additional markers could identify such overlaps. Collectively, this multidimensional analysis provides evidence of a distinct population of responsive TEMRA that is similarly enriched in co-cultures with nAPCs presenting SARS-CoV-2, MMR or Tdap antigens and whose activation depends on IL-15 stimulation.

### Increase in T cells reactive to Tdap and MMR vaccine antigens in uninfected subjects immunized with SARS-CoV-2 vaccine

To determine whether individuals who have received approved SARS-CoV-2 vaccines exhibit an increase in MMR and Tdap specific T cells, we evaluated three uninfected healthy controls before and 2.5 months after they had received the second dose of the Moderna mRNA-based SARS-CoV-2 vaccine. T cell activation was assessed by IFN-γ^+^ ELISpot assays (**Fig. 3A**) and flow cytometry (**Fig. 3B,**C****) as described for Fig. 1D and Fig. 2A,B, respectively. We observed a strong induction of T cells reactive to Spike S1 loaded nAPCs that correlated with a similarly robust activation of T cells reactive to MMR and Tdap antigens that markedly exceeded that observed before vaccination (**Fig. 3A**). No response to nucleocapsid was observed as expected since mRNA-based SARS-CoV-2 encode only for the Spike protein. As observed with SARS-CoV- 2 infected convalescent patients, a high frequency of the IFN-γ^+^ CD4 and CD8 T cells expressed TEMRA markers (**Fig. 3B,**C****).

**Figure 3:**
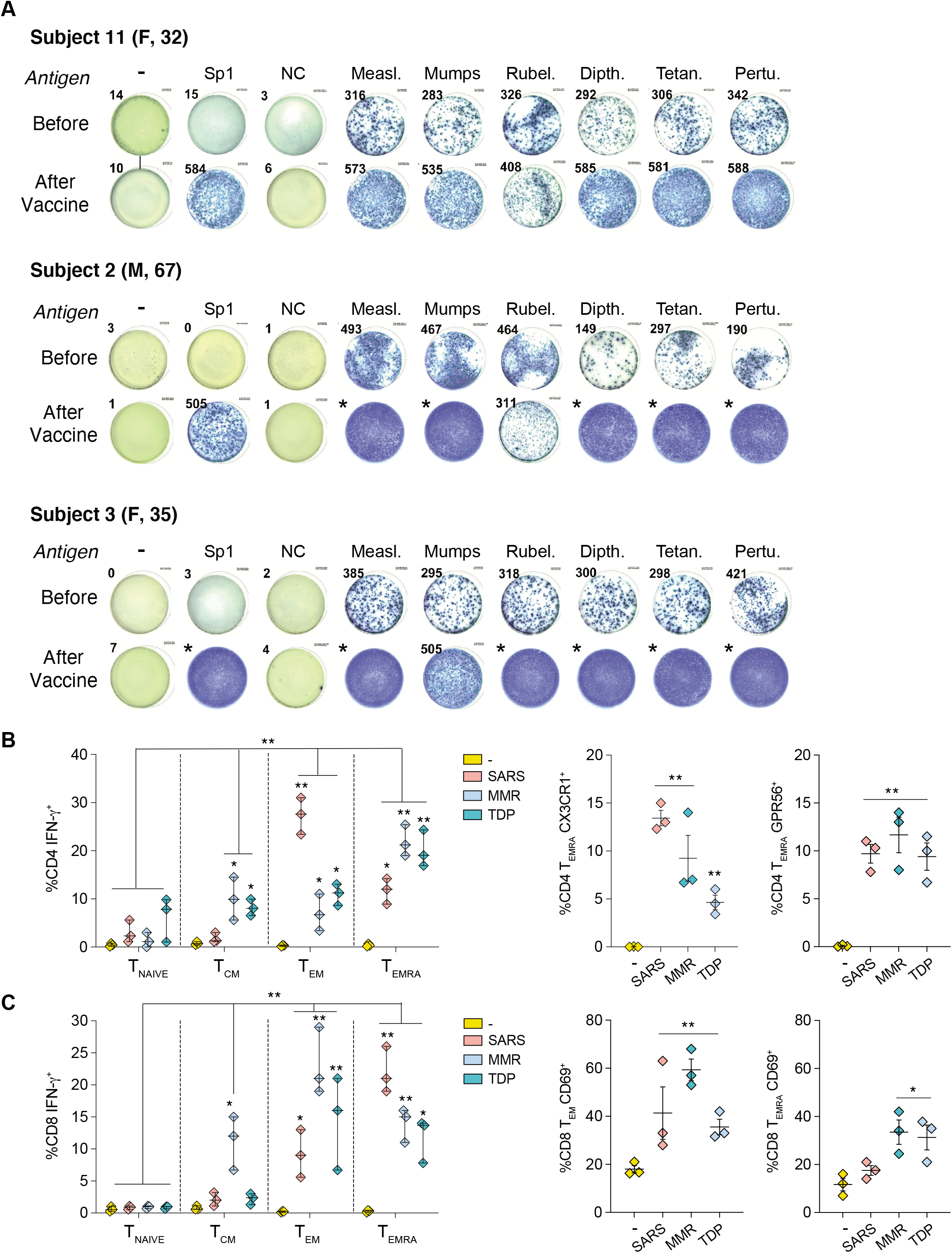
T cell responses to SARS-CoV-2, MMR and Tdap antigens in uninfected, SARS-CoV-2 mRNA vaccinated individuals. **A)** Blood was analyzed from three individuals approximately 3 months before and 2.5 months after receiving the Moderna mRNA-based SARS-CoV-2 vaccine. nAPC generated from blood neutrophils were loaded with individual SARS-CoV-2 and MMR and Tdap antigens and autologous T cell responses were assessed as described in Fig 1A). Images of one well with IFN-γ^+^ spots representative of triplilcates from an ELISpot assay are shown. Asterisk (*) denotes well in which the IFN-γ^+^ spots were too many to be counted. **B-C)** The phenotype of T cells was examined by flow cytometry as in Fig. 2A-B. *p<0.05, **p<0.005 using three-way analysis of variance and Tukey’s multiple comparisons test.

### TCR clonotype and transcriptome of SARS-CoV-2, Tdap and MMR Antigen Reactive T cells

T cell dependent antigen recognition relies on the interaction of TCRs in CD4 and CD8 T cells with peptide loaded MHC class I or II molecules, respectively. TCRα and β chains contain 3 hypervariable loops, termed complementary determining regions (CDR). Of the 3, CDR3 sequences are unique for each clone and are the main contributors to peptide-MHC specificity. Thus, T cells that express the same pair of CDR3 nucleotide sequences are highly likely to be derived from the same clonally expanded T cell. To identify and characterize cells with cross-reactive TCR clonotypes we performed single cell RNA sequencing with T-cell receptor sequence V(D)J capture. We profiled 3 replicate batches of T-cells from a COVID-19-convalescent patient that were exposed to SARS-CoV-2, MMR or Tdap antigen loaded nAPCs, and T cells from a healthy control exposed to SARS-CoV-2 antigen loaded nAPCs, which served as our control for antigen-nonspecific T cell activation. The characteristics of the convalescent patient and healthy control samples are detailed in **Table 1**. After data processing and filtering (see Methods), 15,931 cells remained (range, 833 to 1,663 per sample). Principal component analysis was used to reduce the dimensionality of the dataset for graph based clustering and uniform manifold approximation and projection (UMAP) visualization. This resulted in two large groups of T-cells, one from the healthy controls and the other from the convalescent COVID-19 patients (**Fig. 4A, left panel**), suggesting that the major source of variation in gene expression was between antigen-nonspecific and antigen-specific T cell activation. Accordingly, IL-2RA (CD25), a canonical CD4 and CD8 T cell activation marker that encodes for the IL-2 receptor (*47*) was upregulated only in antigen stimulated (COVID convalescent) T cells (**Fig. 4A, right panel**).

**Figure 4:**
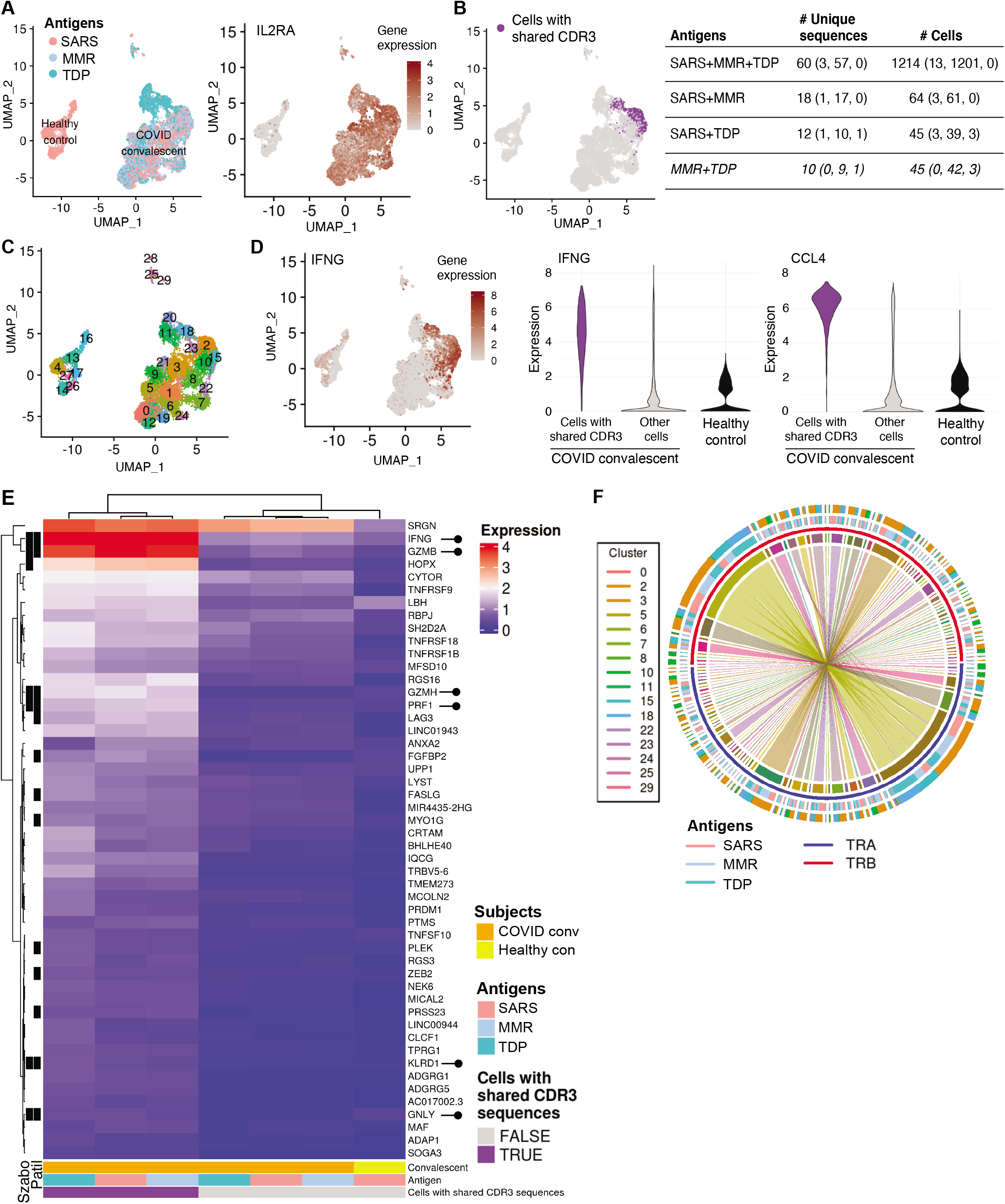
TCR clonotyping and single cell RNA sequencing of T cells. **A**) Uniform manifold approximation and projection (UMAP) plots of single cell gene expression data colored by the antigen used (*left*) and IL-2RA gene expression (*right*). **B)** UMAP plot highlighting cells with shared CDR3 sequences *(left*). Number of clonotypes and cells across indicated antigens SARS-CoV-2 (SARS), MMR and Tdap (TDP) antigens overall and by each COVID-19 convalescent individual (in parentheses) is given. **C)** UMAP plot displaying the clustering scheme. **D)** UMAP plot of single cell gene expression data for IFNG (*left*) and violin plots showing the distribution of expression of IFNG (*center*) and CCL4 (*right*). **E)** Heatmap of mean expression of top 50 marker genes for cells with shared CDR3 sequences by COVID convalescent status and antigen. The black bars on the left-hand side indicate if a gene was identified as a TEMRA marker in Patil et al. 2018 (*50*) (column labelled “Patil”) or in Szabo et al. 2019 (*47*) (column labelled “Szabo”). Rows and columns were grouped by overall expression pattern using hierarchical clustering. Lollipops (*right*) highlight cytotoxic T cell effector molecule genes. **F)** Circos plot of sequences with shared CDR3 sequences. Each sector on the innermost track corresponds to an α chain (TRA, blue) or β chain (TRB, red). The width of a sector is proportional to the number of times the α or β chain sequence occurs. An arc between an α and β chain indicates these sequences are combined in a single CDR3 sequence. The antigen is shown in the third track. The Seurat cluster is shown in the fourth track; not all clusters have cells with shared CDR3 sequences so fewer than 30 clusters are shown. The preponderance of straight lines spanning across the plot shows that in nearly all cases, a given α chain is combined with only one β chain, and vice versa.

Next, we determined whether T cells responsive to SARS-CoV-2 in COVID-19 convalescent patients express the same clonotype CDR3 sequences as T cells responsive to MMR and Tdap. Analysis of the TCR clonotype sequences revealed 12,613 unique clonotypes of which 90 clonotypes met the expected criteria (see Methods) **(Table S1)**. These 90 clonotypes (0.7% of the original 12,613 clonotypes) were expressed in 1,323 cells (8.3% of the cells profiled) (**Fig. 4B**). Each clonotype was unique to the replicate batch and all three replicate batches had cells with shared CDR3 sequences, however they were most concentrated in replicate 2 with 84 of the 90 clonotypes in 1,301 cells (**Fig. 4B**). Reasons for this variance may include permissive HLA haplotype for presentation of peptides that are cross-reactive or the differential retention of cross-reactive effector memory T cells in blood versus the tissue. The majority (60 of 90) of clonotypes occurred in all three antigen samples (SARS-CoV-2, MMR, Tdap), with the remaining 30 present in SARS-CoV-2 plus MMR and SARS-CoV-2 plus Tdap samples (**Fig. 4B**). In addition to the 90 candidate heterologous COVID- 19 CDR3 clonotypes, 10 additional clonotypes overlapped between MMR and Tdap (**Fig. 4B**). Given our interest in the gene expression profile of these specific cells, using graph-based clustering we chose a partitioning of cells that most closely encapsulated the cells containing shared CDR3 sequences. This resulted in a total of 30 cell populations (clusters), where clusters 2, 15, 18 (**Fig. 4C**) were enriched for shared CDR3 sequences (69%, 59%, 54% of cell population respectively). Twelve other clusters contained a small (≤ 15% of cell population) number of cells with shared CDR3 sequences (**Table S1**).

T cells produce a heterogeneous repertoire of effector molecules depending on the subset of T cells that is responding (*26, 48, 49*). IFN-γ serves as a robust marker of a fraction of effector memory T cells following direct TCR activation with message correlating directly with protein (*26*). We found that IFNG overlapped with cells identified with shared CDR3 sequences in UMAP space, with differential expression analyses showing that the levels of IFNG is significantly higher in these cells compared to other T cells in the convalescent group or healthy controls. Likewise, the expression of the chemokine CCL4, also induced by TCR activation of memory T cells (*47, 49*), was significantly higher in the cells with shared CDR3 (**Fig. 4D and Table S2**).

We next sought to identify a gene expression signature for cells that were highly enriched for shared CDR3 sequences (clusters 2, 15, and 18). These clusters contained a total of 1810 cells, of which 1,154 (63.7%) were cells with shared CDR3 sequences (84 of the 90 identified clonotypes) indicating cells with shared CDR3 sequences also have similar gene expression profiles. Differential expression analysis of these clusters relative to all other clusters identified 386 genes that were expressed at significantly higher levels **(Table S2)**. The top 50 of these genes for cells with shared CDR3 sequences (by statistical significance) are shown in a heatmap (**Fig. 4E)**. Of note, of the top 50 genes, 13 were identified as TEMRA markers in Patil et al 2018 (*50*), and 6 were identified as TEMRA markers in Szabo et al 2019 (*47*) (**Fig. 4E, Table S2**)

Further investigation of TCRα and β chain combinations within the shared CDR3 sequences shows they are mostly unique pairings, with only one β chain pairing with 2 α chains (represented by the ‘curved’ arcs) (**Fig. 4F and Table S1**).

### Effect of Tdap and MMR Immunization on COVID-19 disease outcomes

There is growing epidemiological evidence that vaccinations can impact morbidity and mortality beyond their effect on the diseases they prevent (*20, 51*). In COVID-19, a significant association between MMR vaccination status and lower COVID-19 disease severity was observed (*52*) and high titers of mumps antibodies were more likely to be associated with asymptomatic or less severe COVID-19 disease (*53*). These studies were however limited by small sample size (*52, 53*) or survey research methodology (*52*) and may be confounded by co-variables that influence both the likelihood of getting vaccinated with MMR or Tdap and the risk of progressing to severe COVID. To address these challenges, we performed a retrospective cohort study with overlap propensity score weighting at the Cleveland Clinic Health System in Ohio and Florida. All patients tested for COVID-19 between March 8, 2020, and March 31, 2021 were included (73,582 COVID positive patients). The cohort included fewer patients vaccinated with MMR (11,483) than Tdap (36,793) (**Table S3**), consistent with vaccination scheduling, as the single MMR is given at childhood while the Tdap is given as a booster every 10 years and the trivalent MMR vaccine was only available in 1971 (*54*). After adjusting for 44 patient characteristics **(Table 2)**, the two primary endpoints reflecting disease severity (COVID-related hospitalization, and COVID-related admission to the intensive care unit or death) were decreased in patients previously vaccinated for MMR by 38% and 32%, respectively, and in patients previously vaccinated for Tdap by 23% and 20%, respectively **(Fig. 5, Table 2)**. The time interval from vaccination (either MMR or Tdap) to positive COVID-19 test was not significantly associated with outcome, possibly because this cohort is dominated by individuals who had MMR or Tdap vaccines within the past 20 years. Thus, this may not be the ideal dataset to test the effect of interval from vaccination to disease.

**Figure 5:**
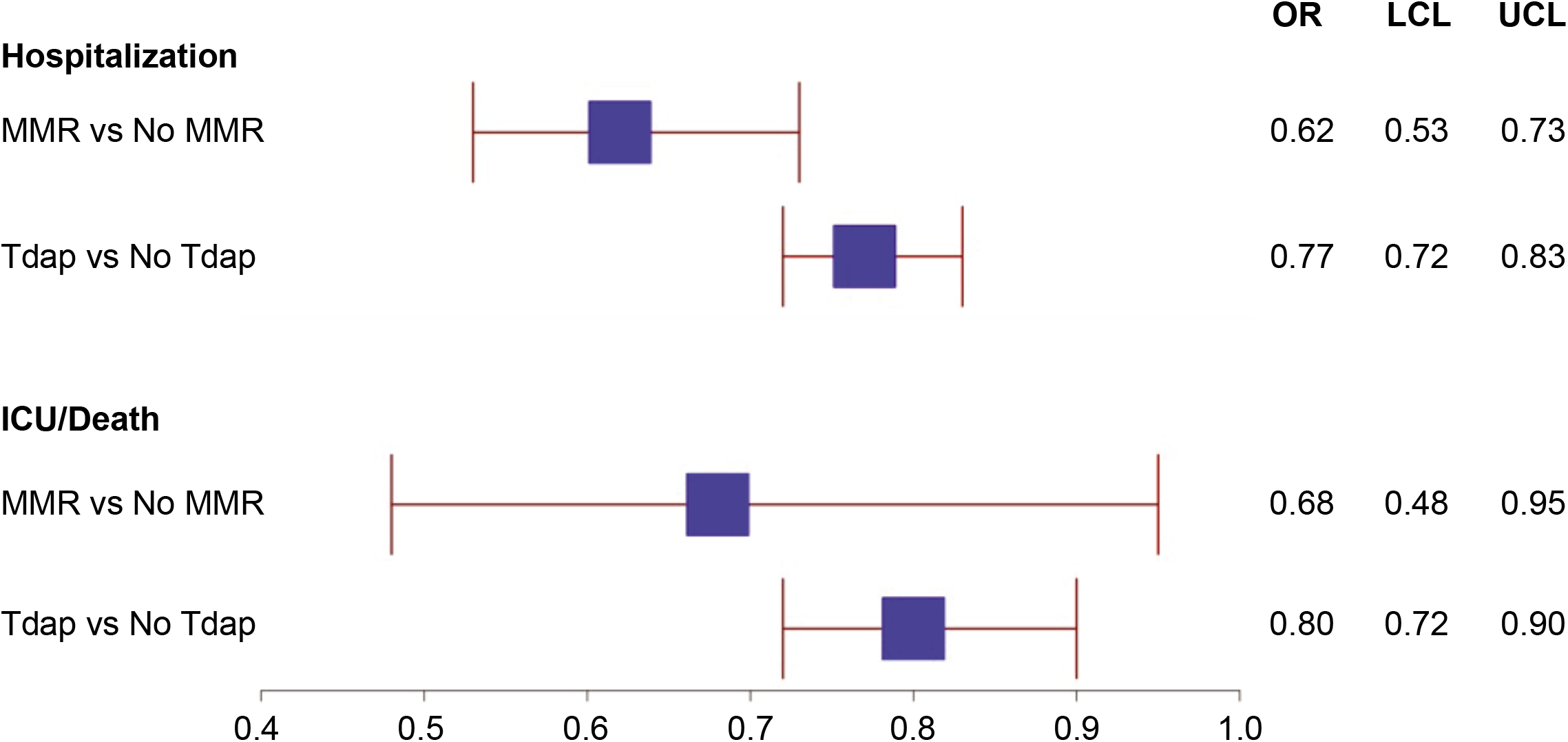
Association of MMR and Tdap vaccination history with hospitalization and progression to ICU stay or death from SARS-cov-2 infection. Overlap Propensity Score Weighted–Analysis Odds Ratios with 95% Confidence Intervals. The upper limit of the 95% confidence intervals for the adjusted odds ratios was less than 1 for both risk of hospitalization and risk of transfer to the intensive care unit or death for patients with a history of prior vaccination for either MMR or Tdap.

**Table 2:**
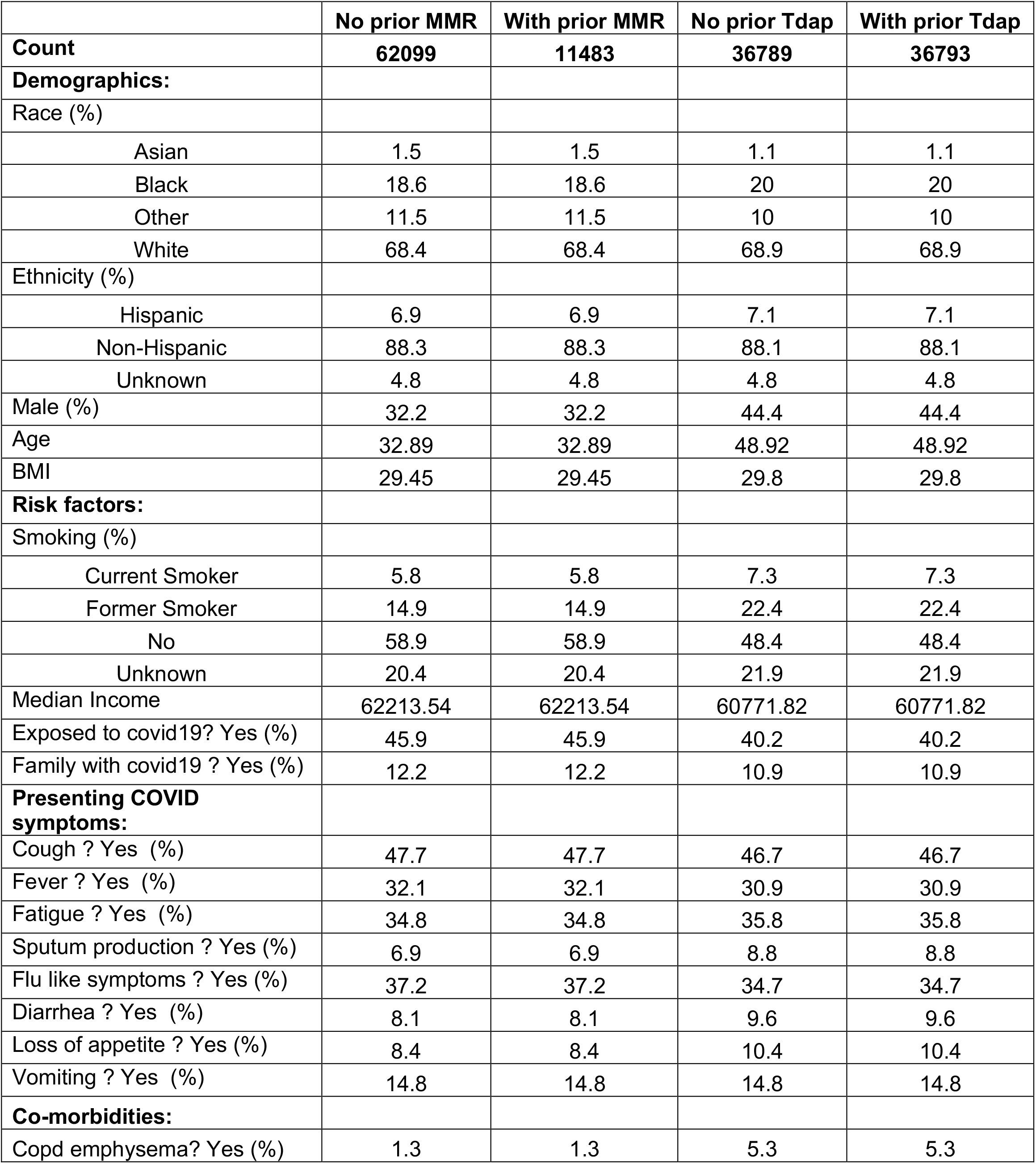
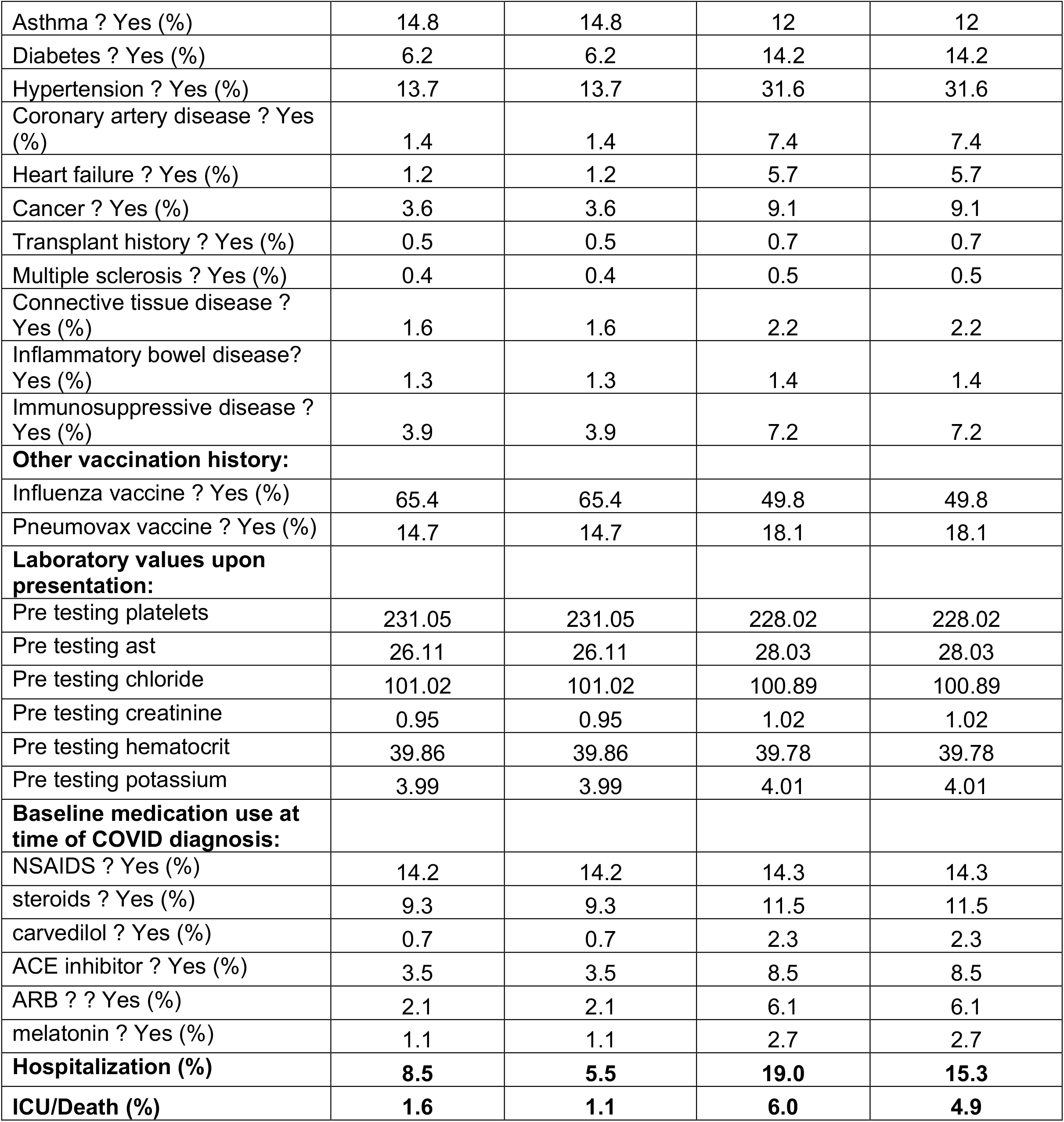
Overlap propensity score–weighted characteristics and disease severity markers (hospitalization, ICU admission/death) among MMR and Tdap vaccination status groups in all COVID positive patients. Our propensity score matching (*69*) was effective at making the two groups comparable as evidenced by the identical scores across patient groups for both the MMR and Tdap comparisons.

|  | No prior MMR | With prior MMR | No prior Tdap | With prior Tdap |
| --- | --- | --- | --- | --- |
| **Count** | **62099** | **11483** | **36789** | **36793** |
| **Demographics:** |  |  |  |  |
| Race (%) |  |  |  |  |
| Asian | 1.5 | 1.5 | 1.1 | 1.1 |
| Black | 18.6 | 18.6 | 20 | 20 |
| Other | 11.5 | 11.5 | 10 | 10 |
| White | 68.4 | 68.4 | 68.9 | 68.9 |
| Ethnicity (%) |  |  |  |  |
| Hispanic | 6.9 | 6.9 | 7.1 | 7.1 |
| Non-Hispanic | 88.3 | 88.3 | 88.1 | 88.1 |
| Unknown | 4.8 | 4.8 | 4.8 | 4.8 |
| Male (%) | 32.2 | 32.2 | 44.4 | 44.4 |
| Age | 32.89 | 32.89 | 48.92 | 48.92 |
| BMI | 29.45 | 29.45 | 29.8 | 29.8 |
| **Risk factors:** |  |  |  |  |
| Smoking (%) |  |  |  |  |
| Current Smoker | 5.8 | 5.8 | 7.3 | 7.3 |
| Former Smoker | 14.9 | 14.9 | 22.4 | 22.4 |
| No | 58.9 | 58.9 | 48.4 | 48.4 |
| Unknown | 20.4 | 20.4 | 21.9 | 21.9 |
| Median Income | 62213.54 | 62213.54 | 60771.82 | 60771.82 |
| Exposed to covid19? Yes (%) | 45.9 | 45.9 | 40.2 | 40.2 |
| Family with covid19 ? Yes (%) | 12.2 | 12.2 | 10.9 | 10.9 |
| **Presenting COVID symptoms:** |  |  |  |  |
| Cough ? Yes (%) | 47.7 | 47.7 | 46.7 | 46.7 |
| Fever ? Yes (%) | 32.1 | 32.1 | 30.9 | 30.9 |
| Fatigue ? Yes (%) | 34.8 | 34.8 | 35.8 | 35.8 |
| Sputum production ? Yes (%) | 6.9 | 6.9 | 8.8 | 8.8 |
| Flu like symptoms ? Yes (%) | 37.2 | 37.2 | 34.7 | 34.7 |
| Diarrhea ? Yes (%) | 8.1 | 8.1 | 9.6 | 9.6 |
| Loss of appetite ? Yes (%) | 8.4 | 8.4 | 10.4 | 10.4 |
| Vomiting ? Yes (%) | 14.8 | 14.8 | 14.8 | 14.8 |
| **Co-morbidities:** |  |  |  |  |
| Copd emphysema? Yes (%) | 1.3 | 1.3 | 5.3 | 5.3 |
| Asthma ? Yes (%) | 14.8 | 14.8 | 12 | 12 |
| Diabetes ? Yes (%) | 6.2 | 6.2 | 14.2 | 14.2 |
| Hypertension ? Yes (%) | 13.7 | 13.7 | 31.6 | 31.6 |
| Coronary artery disease ? Yes (%) | 1.4 | 1.4 | 7.4 | 7.4 |
| Heart failure ? Yes (%) | 1.2 | 1.2 | 5.7 | 5.7 |
| Cancer ? Yes (%) | 3.6 | 3.6 | 9.1 | 9.1 |
| Transplant history ? Yes (%) | 0.5 | 0.5 | 0.7 | 0.7 |
| Multiple sclerosis ? Yes (%) | 0.4 | 0.4 | 0.5 | 0.5 |
| Connective tissue disease ? Yes (%) | 1.6 | 1.6 | 2.2 | 2.2 |
| Inflammatory bowel disease? Yes (%) | 1.3 | 1.3 | 1.4 | 1.4 |
| Immunosuppressive disease ? Yes (%) | 3.9 | 3.9 | 7.2 | 7.2 |
| **Other vaccination history:** |  |  |  |  |
| Influenza vaccine ? Yes (%) | 65.4 | 65.4 | 49.8 | 49.8 |
| Pneumovax vaccine ? Yes (%) | 14.7 | 14.7 | 18.1 | 18.1 |
| **Laboratory values upon presentation:** |  |  |  |  |
| Pre testing platelets | 231.05 | 231.05 | 228.02 | 228.02 |
| Pre testing ast | 26.11 | 26.11 | 28.03 | 28.03 |
| Pre testing chloride | 101.02 | 101.02 | 100.89 | 100.89 |
| Pre testing creatinine | 0.95 | 0.95 | 1.02 | 1.02 |
| Pre testing hematocrit | 39.86 | 39.86 | 39.78 | 39.78 |
| Pre testing potassium | 3.99 | 3.99 | 4.01 | 4.01 |
| **Baseline medication use at time of COVID diagnosis:** |  |  |  |  |
| NSAIDS ? Yes (%) | 14.2 | 14.2 | 14.3 | 14.3 |
| steroids ? Yes (%) | 9.3 | 9.3 | 11.5 | 11.5 |
| carvedilol ? Yes (%) | 0.7 | 0.7 | 2.3 | 2.3 |
| ACE inhibitor ? Yes (%) | 3.5 | 3.5 | 8.5 | 8.5 |
| ARB ? ? Yes (%) | 2.1 | 2.1 | 6.1 | 6.1 |
| melatonin ? Yes (%) | 1.1 | 1.1 | 2.7 | 2.7 |
| **Hospitalization (%)** | **8.5** | **5.5** | **19.0** | **15.3** |
| **ICU/Death (%)** | **1.6** | **1.1** | **6.0** | **4.9** |

## DISCUSSION

Our findings provide definitive cellular and molecular evidence that heterologous adaptive immunity exists between SARS-CoV-2 and antigens present in Tdap and MMR vaccines. We observe enhanced *in vitro* T cell recall responses to Tdap and MMR antigens in individuals with a history of recent SARS-CoV-2 infection or uninfected individuals immunized with the SARS-CoV-2 vaccine, and a strong correlation between the magnitude of the effector memory T cell response activated by exposure to APCs loaded with SARS-CoV-2 antigens and Tdap or MMR vaccine antigens. We identify identical TCR clonotypes in IFNG expressing T cells following activation by SARS-CoV-2, Tdap and MMR antigens, thus providing clear molecular evidence for heterologous immunity. We also demonstrate that the cross-reactive T cells resemble cytotoxic TEMRA, known to contribute to anti-viral immunity. Heterologous immunity can variously alter disease outcomes by providing enhanced immunity or by exacerbating immunopathology or lessening viral control. Our propensity-weighted analysis of a large COVID-19 cohort adjusted for multiple patient characteristics revealed that severe disease outcomes were reduced in MMR or Tdap vaccinated individuals.

Heterologous immunity has been documented in mouse models with examples of protection against widely divergent pathogens (*55, 56*). Innate heterologous immunity can result from antigen-nonspecific functional epigenetic rewiring of monocyte/macrophage precursors in response to one pathogen that induces immunity to a second unrelated infection (*55*). This has been best studied in humans in the context of Bacille Calmette-Guérin vaccine for tuberculosis, which may afford some protection against SARS-CoV-2 (*57-59*). Adaptive heterologous immunity is mediated by memory T cells and antibodies and has been documented for infections by or immunization against bacteria and viruses in experimental mouse models and in humans (*20, 60*). Our data provide direct molecular evidence of overlapping TCRs in T cell clones that respond to SARS-CoV-2 proteins and Tdap and MMR vaccine antigens. The high frequency of overlap in TCR CDR3 sequences across viral (MMR) and bacterial (Tdap) antigens In SARS-CoV-2 infected individuals suggests that heterologous immunity is prevalent in humans. This is consistent with the estimate that effective immunity during a human’s lifetime requires that each of the unique 10^7^-10^8^ TCRs recognize up to 10^6^ theoretical peptides (*62-64*). It is also consistent with studies by Su et al., 2013 (*60*), which detected large numbers of CD4 T cells with a memory-phenotype that recognize viral peptides in unexposed adults. We found that the phenotype and transcriptional profile of the cross-reactive T cell clones primarily comprise activated TEMRA, a cytotoxic effector memory T cell subset unique to humans that promotes anti-viral immunity by eliminating infected cells. For example, the frequency of CD4^+^ TEMRA in Dengue virus correlates with vaccine-elicted protection (*39, 41, 42*) and CD8^+^ TEMRA may control viral load during early HIV infection (*43*). Our ability to detect cross-reactive T cells was enabled by three features of our approach: 1) read out on IFNγ generation, which predominantly identifies activated effector memory versus naïve T cells (*26, 27*); 2) use of highly immunogenic nAPC that generate IL-15, which sensitizes T cells with low TCR binding affinity for antigens (*33-37, 61*); and 3) an epitope unbiased approach in which the relevant peptide epitopes were generated by physiological antigen-processing rather than exposure to a limited set of viral peptide epitopes (*7-10*), which may not represent the specificities of cross-reactive clones generated during a natural SARS-CoV-2 infection To date, it has been difficult to evaluate T cell responses to vaccines due to the lack of sensitive *in vitro* assays. Our approach, described herein, may help bridge this gap.

Prior epidemiological studies show that human childhood Tdap and MMR vaccines reduce the incidence of other infections (*65*), and in the case of MMR, may lower COVID-19 disease severity (*52, 53, 66, 67*). These small observational studies generated interesting correlations but were significantly underpowered and failed to adjust for confounders, limiting the ability to draw meaningful conclusions. Here, we used a large, deeply-phenotyped COVID-19 cohort of close to 74,000 patients, with >300 datapoints per patient, and adjusted for 44 patient characteristics that include demographics, social determinants of health, symptoms upon initial presentation with COVID, co-morbidities, additional vaccination history (influenza and pneumococcal vaccination), laboratory values, and home medication use prior to COVID. Our propensity-weighted methodology allows us to isolate differences attributed to MMR and Tdap vaccination and revealed that severe disease outcomes (hospitalization, transfer to intensive care unit, or death) were reduced in MMR or Tdap vaccinated individuals by 32-38% and 20-23% respectively.

Although there is evidence that SARS-CoV-2 specific memory T cells are generated during natural infection and following vaccination, not enough time has lapsed since the start of the pandemic to assess the longevity (half-life) of these memory T cells (*7, 68*). Effective T cell memory is essential for productive anti-viral immunity particularly to pathogens that evolve to evade recognition by neutralizing antibodies (*1*). This is especially relevant for emerging SARS-CoV-2 variants in the Spike protein, the targets of antibodies that in turn may provide selective pressure for viral escape as population immunity expands. In natural SARS-CoV- 2 infection, the more invariant nucleocapsid protein contains the highest density of SARS-CoV-2 epitopes recognized by CD8 T cells (*14*). In our studies, the robust correlation of MMR and Tdap reactive memory T cells to Spike-S1 and the nucleocapsid protein predicts that these T cells will enhance immunity to SARS-CoV-2 Spike variants, which may emerge as neutralizing antibodies to Spike S1 provide selective pressure for viral escape as population immunity expands. In uninfected individuals immunized with the SARS-CoV-2 vaccine, the markedly enhanced T cell responses to MMR and Tdap antigens compared to the same evaluated prior to vaccination, suggests that protective heterologous T cell immunity in vaccinated individuals may be induced by MMR and Tdap vaccine antigens. Indeed, the observed prevalence of heterologous immunity in our studies may have implications for vaccine development against future novel pathogens, as the effectiveness of vaccines may correlate with their ability to harness pre-existing memory T cells generated by prior infections or vaccines. We posit that intentional MMR or Tdap vaccine-induced heterologous immunity to SARS-CoV-2 could enhance the effectiveness of SARS-CoV-2 vaccines by generating an expanded population of SARS-CoV-2-specific memory T cells (and, in turn, B cells) that respond vigorously to the vaccines and, in countries where SARS-CoV-2 vaccines are not yet available, provide protection from severe disease. The high frequency of T cells reactive to MMR and Tdap in SARS-CoV-2 vaccinated individuals together with the correlation with decreased disease severity independent of age in our retrospective analysis of COVID-19 patients suggests that MMR or Tdap vaccination may enhance the efficacy and durability of SARS-CoV-2 vaccines in individuals of all ages.

In conclusion, our studies provide evidence of broad cross-reactivity between T cells responsive to SARS-CoV-2, MMR and Tdap antigens in humans. The breadth of cross-reactive CDR3 sequences to these three distinct pathogens suggests that adaptive heterologous immunity is prevalent in humans. The correlation of MMR and Tdap vaccination in COVID-19 patients with a decrease in disease severity may reflect the observed cross-reactivity of MMR and Tdap vaccine antigens not only with Spike-S1 protein but also with the relatively invariant nucleocapsid protein. A signature of cytotoxic effector memory CD4 T cells (TEMRA) in responding cross-reactive cells, and the reported prevalence of this T cell subset in recovered COVID-19 patients (*44*), suggests that this T cell subset may promote robust, heterologous anti-viral immunity in COVID- 19. Finally, our studies predict that MMR or Tdap vaccination together with approved SARS-CoV-2 vaccines that currently target Spike-S1 protein may afford greater protection, particularly against emerging Spike variants, than COVID-19 vaccines alone.

## ACKNOWLEDGEMENTS

This work was supported by National Institutes of Health R01HL065095 (T.M.), R01AI152522 (T.M.), R01NS097719 (L.J.) and a generous donation from Barbara and Amos Hostetter and the Chleck Foundation (D.R.W.). We would like to thank Brigham and Women’s Hospital and MassCPR for their support in collecting samples from SARS-CoV-2 infected patients, Colin Heyson for statistical advice, Pascal Yazbeck and Chi-An Cheng for technical assistance, and Jon Aster for scientific discussion and advice.

## AUTHOR CONTRIBUTIONS

V.M and X.C. performed, and V.M. designed and analyzed all cell-based studies; T.G. performed studies on plasma samples; M.L.S. and B.D.J. completed the scRNA sequencing and TCR clonotyping analysis; M.D. and L.R.B. collected and managed clinical blood samples from COVID-19 participants; X.J. participated in data analysis; M.W.K and L.J. participated in study design, data interpretation and data analysis; A.H.L. provided valuable advice; T.N.M. conceptualized the research and designed and supervised the experiments; T.M., V.M., X.C., A.H.L., M.L.S., M.W.K. and L.J. wrote the manuscript. All authors edited the manuscript.

## METHODS

### Ethics statement

All procedures performed were in accordance with ethical standards. Blood samples were obtained from consented healthy, self-reporting SARS-CoV-2 uninfected volunteers under a Mass General Brigham Institutional Review Board (IRB)-approved protocol (1999P001694). Patients signed informed consent to participate in a Mass General Brigham IRB-approved COVID-19 observational sample collection protocol (2020P000849).

### Serum immunoglobulin Simoa Assays

SARS-CoV-2 serological Simoa assays for IgG against four viral antigen S1, Spike, Nucleocapsid, and RBD were prepared and preformed as previously described (*31*). Briefly, plasma samples were diluted 4000-fold in Homebrew Detector/Sample Diluent (Quanterix Corp.). Four antigen-conjugated capture beads were mixed and diluted in Bead Diluent, with a total of 500,000 beads per reaction (125,000 of each bead type). Biotinylated anti-human IgG antibodies (Bethyl Laboratories A80-148B) were diluted in Homebrew Detector/Sample Diluent to final concentrations of 7.73ng/mL. Streptavidin-β-galactosidase (SβG) was diluted to 30 pM in SβG Diluent (Quanterix). The serology assay was performed on an HD-X Analyzer (Quanterix) in an automated three-step assay. Average Enzyme per Bead (AEB) values were calculated by the HD-X Analyzer software. All samples were measured in duplicates.

### Plasma cytokine assays

Plasma cytokines were measured in plasma samples using the CorPlex Cytokine Panel (Quanterix Corp), which included sample diluent buffer. Plasma samples were diluted 4-fold in sample diluent buffer and assays were performed following the CorPlex manufacturer protocols. Each CorPlex cytokine panel kit was analyzed by the SP-X Imaging and Analysis System (Quanterix Corp.). All samples were measured in duplicates.

### Cell isolation, treatment and culture

#### Blood and serum collection

Peripheral blood was drawn into tubes containing trisodium citrate, citric acid and dextrose (Vacutainer ACD Solution A, BD). Serum was obtained by drawing blood into BD Vacutainer™ Venous Blood Collection Tubes SST, followed by centrifugation at 2500xg for 30 min and removal of the resulting supernatant.

#### Generation of complexed anti-FcγRIIIB (AAC)

Anti-FcγRIIIB (3G8) (Biolegend) was conjugated to FITC-Ovalbumin (#O23020, Thermofisher) by Biolegend as a custom order and referred to as antibody-antigen conjugate (AAC). Importantly, Ovalbumin in the AAC served as a model antigen in mouse models (*25*) but is irrelevant for our human studies.

#### Human blood treatments to generate neutrophil derived APC (nAPC)

10mls human blood was supplemented with GM-CSF (10 ng/ml) for 30 min at 37C followed by addition of 30µg AAC or FITC-IgG isotype control for 2 hrs at 37C. Blood was then incubated with Hetasep (STEMCELL Technologies) according to manufacturer protocols to deplete red blood cells and enrich leukocytes. Neutrophils were isolated from the leukocyte-rich plasma layer using a Easysep Neutrophil enrichment kit (STEMCELL Technologies) and placed in RPMI media, which was supplemented with 10% autologous serum, penicillin/streptomycin (50 U/ml penicillin and 50 mg/ml streptomycin) and 20ng/ml GM-CSF. After 48 hours, cells were harvested using Accutase and evaluated by flow cytometry for surface markers of APCs.

#### Monocyte isolation and culture to generate monocyte-derived DCs (moDC)

Peripheral blood mononuclear cells were isolated using Lymphoprep (Stemcell technologies, Canada). Monocytes were positively selected by anti-CD14-coated magnetic beads (Miltenyi biotec, Germany) to >98% purity. Monocytes were cultured in complete medium supplemented with GM-CSF (50ng/ml) and IL-4 (10ng/ml). Cells were harvested after 7 days and evaluated for surface markers CD11c, HLA-DR and CD14.

#### T cell isolation

PBMCs were isolated from peripheral blood using Lymphoprep (Stemcell technologies, Vancouver, Canada) density gradient medium, aliquoted in 1ml cryopreservation tubes at a concentration of 5 million cells/ml and frozen. The tubes were thawed after 2 days to isolate CD3 T cells for co-culture studies. For isolation of CD3+ T cells, negative selection was performed using EasySep Human T cell isolation kit (#17951, Stemcell technologies, Vancouver, Canada). The CD3+ T cells were labelled with 1µM Cell Trace Violet dye (#C34557, ThermoFisher Sientific) according to manufacturer’s instructions just prior to setting up their co-cultures.

#### Co-culturing nAPC/moDC and T cells

nAPCs derived from neutrophils and moDCs were harvested and co-cultured with Cell trace Violet labelled CD3+ T cells isolated from PBMCs at a ratio of 1:5 (nAPC:T cells) on a IFNγ ELISpot plate and incubated for 18h. Additionally, the co-cultures were incubated with the following antigens individually or in combinations indicated in the Result section: SARS-COV-2 antigens 5µg/ml Spike-S1 subunit (S1-fc), 8.85µg/ml Spike S2 subunit (S2-his), 7.8µg/ml Spike S1 receptor binding domain (RBD-fc), 2.35µg/ml Nucleocapsid protein (NC his); inactivated Measles, Mumps, Rubella viral preparations at 5µg of total protein/ml; 5µg/ml of heat inactivated Pertussis toxin, Diphtheria toxin and Tetanus toxoid (See Table 2).

#### Antibody blocking treatments

Neutralizing antibodies were used as follows. Anti-IL15(1) (Invitrogen, #16-0157-82) at 5µg/ml, anti-IL15(2) (R&D systems, #MAB247) at 5µg/ml, anti-IL1β (R&D systems, #MAB201) 1.6µg/ml and anti-IL18 (MBL, #D044-3) at 1.8µg/ml. Antibodies remained during the entire period of co-culture.

### Interferon Gamma (IFNγ) ELISpot assay

ELISpot kits to measure the secretion of human IFN-γ (R&D Systems, #EL285) were used according to manufacturer’s protocols. Fresh CD3 T cells (Cell trace violet labelled) isolated from PBMCs and nAPCs were plated in triplicates at a ratio of 1:5 (nAPC:CD3 T cells) per well and incubated with indicated antigens and/or blocking antibodies for 18h. Samples were processed according to manufacturer’s protocol and results were quantitated using an ELISpot reader (CTL ImmunoSpot® S6 Fluorescent Analyzer). Results are a mean of triplicate wells and reported as number of spots per million T cells.

### Cell cytokine detection and analysis

nAPCs and monocyte-derived DCs (moDC) were cultured for 72h and supernatants were collected and analyzed for cytokine and chemokine levels using human cytokine 42-plex discovery assay (Eve Technologies, Calgary, AB).

### Flow cytometry

Flow cytometry was performed on a FACSCanto II. FCS (flow cytometry standard format) 3.0 data file was used to export data that was analyzed using FlowJo (Mac version 10.7). Compensation controls were created for each fluorochrome. BD multicolor compensation beads and cells were used to set up compensation for the individual fluorochromes. For all experiments, cells were stained with the Fixable Viability Dye eFluor 780 (ThermoFisher) to gate out dead cells. Forward and side scatter gates were used to discriminate doublets and debris (FSC-A, FSC-H, SSC-A x SSC-H). Matched isotypes were used as controls and negative gating was based on FMO (fluorescence minus one) strategy. Only viable cells were included for the studies.

For surface staining, single cell suspensions in FACS buffer (PBS supplemented with 2% FCS and 2mM EDTA) were incubated with human TruStain FcX for 20 min at 4C. Samples were incubated with the indicated fluorochrome-conjugated antibodies for 30 min at 4C, washed with PBS and fixed with 1% paraformaldehyde.

To evaluate surface markers on nAPCs, antibodies to the following were used: CD15, CD66b, CD11c, HLA-DR, CD40, CD86 and CCR7 (**Source Table**). Within the viable single cell population, CD11c+ and HLA-DR+ events were further gated for CD66b and CD15 expression. The CD11c+, HLA-DR+, CD15+ and CD66b+ population was further analyzed for CD40, CD86 and CCR7 markers.

To evaluate cell surface markers on T cells, antibodies to the following markers were used: CD3, CD4, CD8, CD45RA, CCR7, CD27, GPR56, CX3CR1, IFN-γ (**Source Table**). Subsets of T cells were classified based on CD45RA, CCR7 and CD27. Live, singlet CD3+ cells were assessed for proliferation by monitoring CFSE stained T cells for Cell trace violet signal dilution. For detection of intracellular IFN-γ, T cells were treated with Brefeldin A (3ug/ml) for 5h, stained with surface markers, fixed for 30min, permeabilized with BD Perm/Wash (BD Biosciences) and stained with anti-IFN-γ (**Source Table**). Cells were washed with permeabilization buffer and analyzed by flow cytometry.

### Flow cytometric data analysis

Sample files were exported as FCS 3.0 files from FACSDiva and imported into flowJo v.10.7.1 software for subsequent analysis. The following plug-ins were used: Downsample (1.1), t-distributed stochastic neighbor embedding (tSNE) and FlowSOM (2.6). To visualize the high-dimensional data in two dimensions, the t-SNE algorithm was applied on data. Cells were selected for each sample at random, downsampled and merged into a single expression matrix prior to tSNE analysis. tSNE was performed unsupervised from a maximum of 5,000 randomly selected cells from each sample, with a perplexity set to 80, using the implementation of tSNE plugin in flowJo. Events were identified by gating on live, singlet intact CD3+CD4+ or CD8+ T cells and were included in generating the tSNE plots. The Barnes-Hut implementation of t-SNE with 1,000 iterations, a perplexity parameter of 30, and a trade-off θ of 0.5, was used for applying the dimensionality reduction algorithm. The output was in the form of 2 columns corresponding to t-SNE dimension 1 and dimension 2. t-SNE maps were generated by plotting each event by its t-SNE dimensions in a dot-plot.

Intensities for markers of interest were overlaid on the dot-plot to show the expression of those markers on different cell islands and facilitate assignment of cell subsets to these islands using FlowSOM plugin. Samples were examined by running tSNE with the following markers: CD4, CD45RA, CCR7, CD27, GPR56, CX3CR1, IFN-γ. Phenotypic characteristics of the cell island are shown as heatmaps.

### Statistical analysis

Statistical analyses for cell-based assays were performed using Graphpad prism 8 (LaJolla, CA), and JMP10 software (SAS Institute, Inc, USA). All the data included in the studies are expressed as mean ± SEM. *P <0.05 and **P <0.005 was considered significant.

**Source Table:** Antigens used to load antigen presenting cells (APCs)

| ANTIGEN | SOURCE | Catalog# |
| --- | --- | --- |
| SARS-CoV-2 S1 protein, Fc-Tag | Acro Biosystems | S1N-C5255 |
| SARS-CoV-2 S protein RBD, Fc-Tag | Acro Biosystems | SPD-C5255 |
| SARS-CoV-2 S2 protein, His-Tag | Acro Biosystems | S2N-C52H |
| SARS-CoV-2 Nucleocapsid protein, His-Tag | Acro Biosystems | NUN-C5227 |
| Rubella Grade III Antigen Protein | Aviva Systems Biology | OPMA04522 |
| Inactivated (gamma radiated) Native Measles Virus | Bio-Rad | PIP013 |
| Inactivated (gamma radiated) Native Mumps Virus | Bio-Rad | PIP014 |
| Pertussis Toxin (heat inactivated before use) | Gibco Invitrogen | PHZ1174 |
| Tetanus Toxoid | EMD Millipore | 582231 |
| Diphtheria Toxin | SIGMA | D0564 |

**Source Table:** Antibodies used for flow cytometry

| Antibody | Company | Clone | Cat# |
| --- | --- | --- | --- |
| CD10 | Ancell Technologies | AHN1.1 | 164-040 |
| CD11c | Biolegend | 3.9 | 301619 |
| HLA-DR | Biolegend | L243 | 307639 |
| CD40 | Biolegend | 5C3 | 334321 |
| CD86 | Biolegend | IT2.2 | 305419 |
| CCR7 | Biolegend | G043H7 | 353214 |
| CD4 | Biolegend | SK3 | 344646 |
| CD8 | Biolegend | SK1 | 344708 |
| CD45RA | Invitrogen | HI100 | 11-0458-42 |
| CX3CR1 | Invitrogen | 2A9-1 | 341612 |
| GPR56 | Biolegend | CG4 | 358206 |
| IFN-γ | Invitrogen | 4S.B3 | 12-7319-42 |
| CD66b | MACS Miltenyi biotech | Tet2 | 130-117-811 |
| CD3 | Biolegend | UCHT1 | 300456 |
| CD27 | BD Biosciences | M-T271 | 560609 |
| CD69 | Biolegend | FN50 | 310938 |

### V(D)J clonotyping and scRNA-seq

#### Sample preparation

nAPCs were co-cultured with CD3+ T cells isolated from PBMCs at a ratio of 1:5 (nAPC : T cells) and incubated for 18h. The co-cultures were incubated with SARS-COV-2 antigens (5µg/ml S1-fc, 7.8µg/ml RBD-fc, 2.35µg/ml NC his) or Measles, Mumps and Rubella viral preparations (5µg of total protein/ml) or 5µg/ml of Diphtheria toxin, Tetanus toxoid and Pertussis toxin was used. After 18h the cells were harvested using Accutase and were aliquoted in 1ml cryopreservation tubes at a concentration of 1.2 million cells/ml and frozen at -80C. Samples were sent to MedGenome, Inc. Foster City, CA, for library preparation and sequencing.

### Library preparation and sequencing

Cryopreserved cells were thawed and washed twice with Dulbecco’s phosphate-buffered saline (Fisher Scientific cat# SH3026402) plus 0.04% bovine serum albumin (Fisher Scientific cat# AM2618) then stained with hashtag antibodies (BioLegend, Cat#s 394661, 394663, 394665, or 394666) according to the manufacturer’s protocol. Cells were counted and checked for viability >80%. Stained cells were brought to ∼1000 cells/ul, pooled equally in sets of four samples, and loaded onto the 10x Chromium instrument (10x Genomics) to generate single-cell gel beads in emulsion. For 5’ multiomics including gene expression, TCR-sequencing, and cell-hashing seq, we used the 10x Single Cell V(D) Kit with Feature Barcoding (10x Genomics, Cat# 1000006) along with the Human V(D) Enrichment Kit (10x Genomics, Cat# 10000005) following the manufacturer’s protocol. Libraries were checked for concentration using Qubit dsDNA HS assay kit (Fisher Scientific, Cat# Q32854) and size distribution using TapeStation (Agilent). Libraries were then sequenced on NovaSeq (Illumina) to a sequencing depth of 50,000 reads per cell for gene expression libraries and 10,000 reads per cell for TCR and hashtag libraries. Cell Ranger reads were aligned to reference GRCh38-2020-A, cellular barcodes were demultiplexed, and unique molecular identifiers and antibody capture were quantified using 10x Genomics’ Cell Ranger software (version 4.0.0).

### Data analysis

scRNA-seq and V(D)J data preprocessing. 10X Genomics Cell Ranger Software Suite (v.4.0.0) (*70*) was used to process raw fastq files to expression tables and assemble V(D)J clonotypes, performing all the necessary barcode processing, mapping to the Human reference genome (GRCh38, GENCODE v32/Ensembl 98) and unique molecular identifier (UMI) expression counting; each batch contained an estimated > 7K cells. The gene-cell matrix of all cells was analyzed in R v4.0.3 with Seurat v3.9.9 (*71*). Individual antibody sample hashtags were used to distinguish cells from the four samples within batch (Normal, Convalescent Covid, MMR and Tdap), requiring at least 100 antibody reads present per cell and tag specificity of >80% to a single sample. V(D)J Clonotypes were linked to the scRNA expression by common cell barcode. The following cell filtering criteria were then applied: gene number greater than 1,000, unique UMI count between 1,000 and 50,000 and mitochondrial gene content < 10%. After filtering, a total of 15,931 cells were left across all 12 samples for the subsequent analysis, of which 14,362 have a V(D)J clonotype defined. All samples were processed together and the matrix was normalized using ‘LogNormalize’ method with default parameters. Then, the top 2,000 variable genes were identified using the ‘vst’ method from Seurat’s FindVariableFeatures function which is used for scaling, dimensionality reduction and clustering. The variables batch, S.Score and G2M.Score from the CellCycleScoring function, percent.mito, and nFeature_RNA were regressed out using ScaleData and PCA was performed. Finally, UMAP and graph-based clustering was performed using the top 50 principal components for visualizing the cells. A cluster resolution of 1.75 was chosen for downstream analysis.

### Identification of candidate heterologous T cells

The dataset contained a total of 12,613 unique clonotypes (alpha and beta chain amino acid sequences) in 15,931 cells. Clonotypes which 1) occurred in a minimum of 3 cells (118 clonotypes); 2) were present in a SARS-CoV-2 sample and at least one of MMR or Tdap sample (92 clonotypes); and 3) and was absent from the healthy control sample (2 clonotypes) were considered. The filter resulted in 90 clonotypes in 1,323 cells highly enriched in a few clusters (clusters 2 – 69% of cells, 10 – 15% of cells, 15 – 59% of cells, 18 – 54% of cells). Batch 2 contained the most heterologous clonotypes with 84, with Batch 1 having 5 clonotypes and Batch 3 just 1.

### Differential gene expression analysis

Wilcoxon rank-sum tests as implemented in Seurat v. 4.0.0 (FindMarkers function) were used to perform differential gene expression (DEG) analysis. For enriched clusters (2, 15,19) containing cells with candidate heterologous T cells (as outlined above), DEGs were generated relative to all of the other clusters. A gene was considered significant with adjusted *P* < 0.05. Only genes upregulated relative to all other cells were considered as markers for the purposes of this analysis.

The heatmap plot was created using the R package ComplexHeatmap, version 2.6.2. The Circos plot was generated using the R package circlize, version 0.4.12.

### Patient selection

The Cleveland Clinic COVID-19 Enterprise Registry was created on March 17, 2020 as a resource for COVID-19 research across the health system. More than 300 data points are extracted from the electronic health record through a combination of manual pulls and validated natural language processing algorithms on all patients tested for COVID-19 in our facilities in Ohio and Florida (*72, 73*) (18 regional hospitals and 220 outpatient locations). A waiver of informed consent (oral or written) from study participants in the COVID- 19 registry was granted by the Cleveland Clinic Health System institutional review board. For this study, we included all COVID positive patients diagnosed between March 2020 and March 31, 2021. Infection with SARS-CoV-2 was confirmed by laboratory testing using the Centers for Disease Control and Prevention reverse transcription–polymerase chain reaction SARS-CoV-2 assay.

### Statistical Analysis

All descriptive statistics were reported as counts (percentages) or median (interquartile ranges [IQRs]). For comparison of demographic variables and comorbidities among cohorts, Wilcoxon signed-rank tests were used for numeric variables, while χ2 or Fisher exact tests were used for categorical variables.

To address differences in baseline characteristics of non-MMR/Tdap vaccinated and MMR/Tdap vaccinated patients, we leveraged appropriate statistical methodology to study our research questions. Overlap propensity score weighting was performed to address potential confounding in comparing non-MMR/Tdap vaccinated and MMR/Tdap vaccinated patients given their baseline differences. The overlap propensity score weighting method was chosen given its benefits of preservation of numbers of individuals in each group and of achieving higher levels of precision in the resulting estimates. This methodology is preferred when the propensity score distributions among the groups are dissimilar and when the propensity scores are clustered near the extremes (i.e., close to zero or one). A propensity score for being MMR/Tdap vaccinated was estimated from a multivariable logistic regression model. For the outcomes of hospital and intensive care unit (ICU) admission or death of COVID-19 test-positive patients, the propensity score logistic regression model included covariates that were found to be associated with the outcome in our previous work.

The overlap propensity score weighting method was then applied where each patient’s statistical weight is the probability of that patient being assigned to the opposite group. Overlap propensity score weighted logistic regression models were used to investigate associations between vaccine status and the probability of hospital admission for COVID-19, and ICU admission or death for COVID-19 illness. The results are thus reported as weighted proportions, odds ratios, and 95% confidence intervals.

To address the effect of the time interval between date of vaccine and date of COVID test to the outcomes, we used the time interval as a covariate into multivariable logistic regression models, adjusting for the same covariates as with the overlap propensity scoring models. The time interval is modeled with restricted cubic splines because of suspected nonlinear effects.

All statistical analyses were performed using R 4.0. P values were 2-sided, with a significance threshold of 0.05.

## SUPPLEMENTAL INFORMATION

**Supplemental Figure S1:**
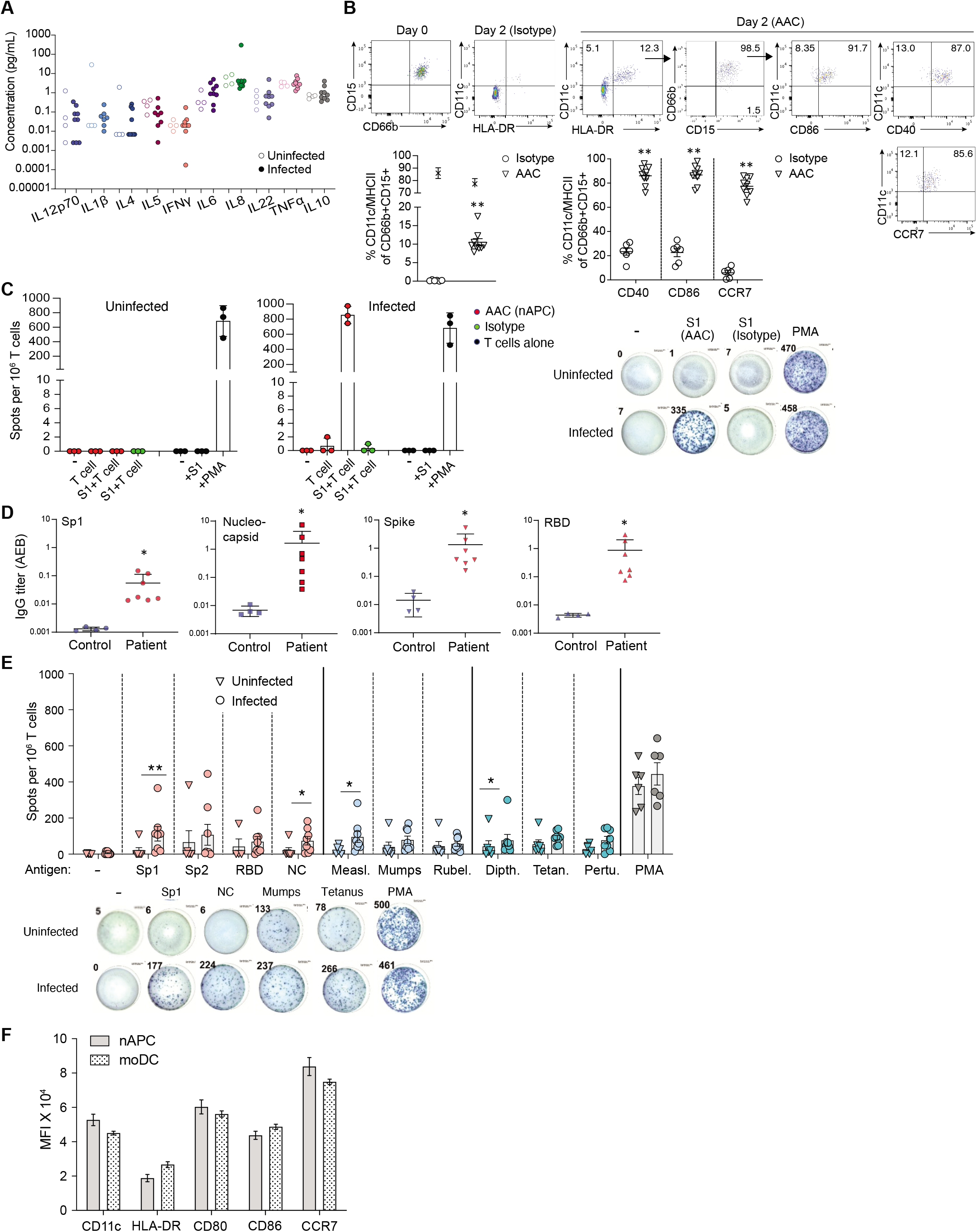
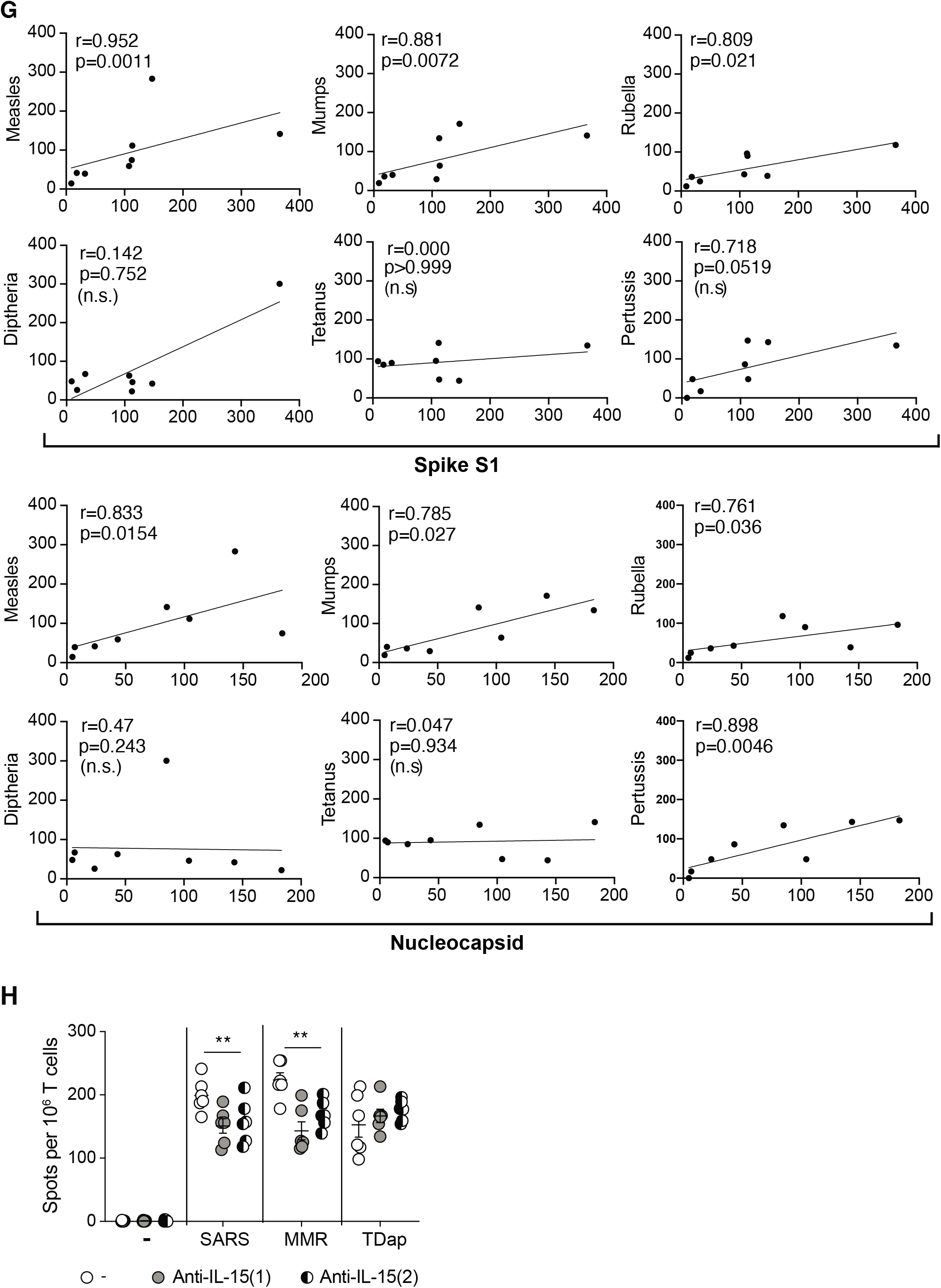
Sera cytokine and antibody profiles, phenotype of generated nAPCs and T cell responses to antigen loaded monocyte derived dendritic cells (moDC). Related to Figure 1. **A)** Cytokine profile in sera of exposed and unexposed individuals. **B)** Representative flow cytometric plots assessing purity of isolated human blood neutrophils. Gating strategy to assess nAPC generation from human neutrophils treated with antibody Isotype and antibody conjugate (AAC) for CD11c+MHCII+ and CD40/CD86/CCR7 on day 2 after culture. Graphs for frequency of cells with indicated markers are shown below. Percent survival was evaluated by exclusion of Fixable Viability Dye positive cells. **p<0.005 using multiple t-tests between paired samples. **C)** Representative ELISpot assay (samples in triplicate) for an uninfected and infected donor highlighting controls used for the assay. Neutrophils incubated with antibody isotype or AAC to generate nAPCs were used in all assays. A ratio of 1:5 (nAPC:T cells) were co-cultured in each well. Representative images of the ELISpot wells are shown. **D)** IgG, IgA and IgM titers in plasma of unexposed and exposed donors were evaluated as in Figure 1B. A significant difference between uninfected and infected individuals was observed for IgG against each of the four viral antigens (Mann Whitney test, p<0.05). **E)** Monocyte derived DCs from a subset of donors were co-cultured with autologous CD3 T cells and analyzed for IFNγ generation using an ELISpot assay as in Figure 1A. *p<0.05; **p<0.005 by two-tailed Mann-Whitney test with Bonferroni correction for multiple comparisons. Representative images of wells with IFN-γ^+^ spots are shown. **F)** Median fluorescence intensity (MFI) for CD11c, HLA-DR and T cell co-stimulatory markers (CD40, CD86) and CCR7, were evaluated on nAPCs and moDCs. No significant differences were observed using multiple t-tests between pair of samples. **G)** Correlations of T cell responses to moDCs pulsed with Spike-S1, nucleocapsid and MMR and Tdap vaccine antigens using Spearman’s rank correlation coefficients are shown between different antigens (*Y axis*) and Spike-S1 or Nucleocapsid (*X axis*). **H)** moDCs co-cultured with autologous CD3 T cells for 18h as in E) were analyzed for IFN-γ^+^ spots in the presence of two independent anti-IL15 blocking antibodies. Two-way analysis of variance and Dunnett’s multiple comparison test was used. **p<0.005.

**Supplemental Figure S2:**
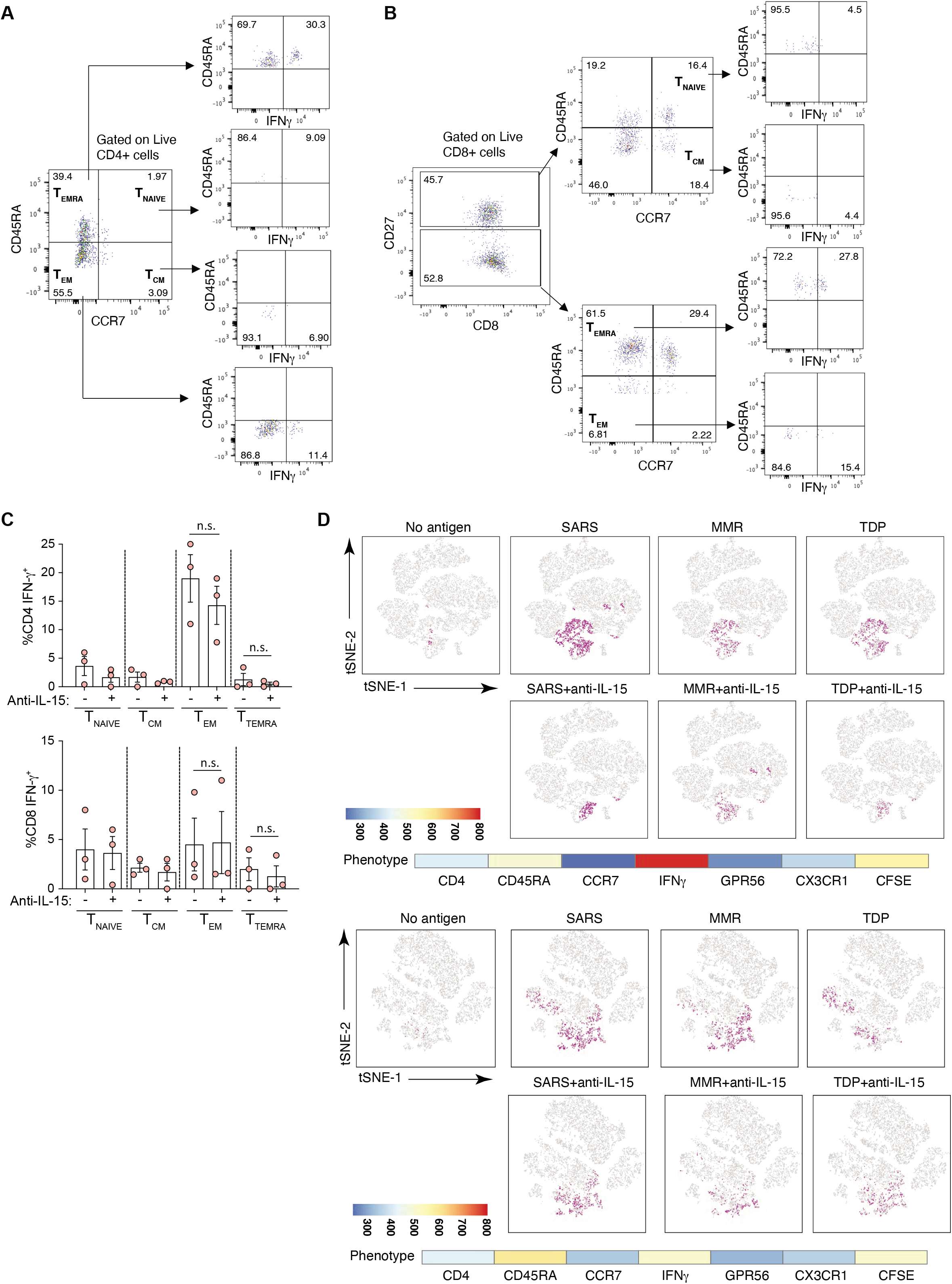
Gating strategies for T cell subsets, phenotype of T cells co-cultured with SARS-CoV-2 antigen loaded monocyte derived dendritic cells (moDC) and unsupervised clustering of T cells co-cultured with antigen loaded nAPC from two infected donors. Related to Figure 2. **A-B)** Gating strategy to assess for naïve, effector and memory populations as shown on representative plots for live CD4 (A) and live CD8 (B). The cells were further gated to show IFN-γ^+^ cells. **C)** T cells co-cultured with moDCs and treated with anti-IL-15 as described in Supplemental Figure 1H) were treated with Brefeldin A for 5h and analyzed by flow cytometry to characterize IFNγ producing CD4 and CD8 T cell subsets. Two-way analysis of variance and Dunnett’s multiple comparison test was used. n.s., not significant. **D)** FLOWSOM based visualization on a tSNE plot. tSNE plots of flow cytometry datasets for CD4+ live T cells analyzed as in Figure 2D for additional two infected donors. The phenotypical evaluation of overlapping population between the antigens are shown along with a continuous scale.

**Supplemental Table S1:** TCRα and β chain combinations. List of the 90 shared CDR3 sequences and cells count presence within sample and cluster.

**Supplemental Table S2:** Differentially expressed genes. List of 386 marker genes identified comparing clusters 2, 15, and 18 against all other clusters. Columns include gene symbol, raw p-value, average log fold change, percent presence within clusters of interest, percent presence outside clusters of interest, adjusted p-value and presence of genes associated with TEMRA cell type identified in Patil et al. 2018 and Szabo et al. 2019.

**Supplemental Table S3:** Characteristics of all COVID-19 patients who were hospitalized in those who were admitted to ICU or died by prior MMR or Tdap vaccination history.

## REFERENCES

1. N. N. Jarjour, D. Masopust, S.C. Jameson, T Cell Memory: Understanding COVID-19. Immunity 54, 14–18 (2021).

2. C. Rydyznski Moderbacher et al., Antigen-Specific Adaptive Immunity to SARS-CoV-2 in Acute COVID-19 and Associations with Age and Disease Severity. Cell 183, 996–1012 e1019 (2020).

3. A. T. Tan et al., Early induction of functional SARS-CoV-2-specific T cells associates with rapid viral clearance and mild disease in COVID-19 patients. Cell Rep 34, 108728 (2021).

4. D. Weiskopf et al., Phenotype and kinetics of SARS-CoV-2-specific T cells in COVID-19 patients with acute respiratory distress syndrome. Sci Immunol 5, (2020).

5. I. Schulien et al., Characterization of pre-existing and induced SARS-CoV-2-specific CD8(+) T cells. Nat Med 27, 78–85 (2021).

6. R. Zhou et al., Acute SARS-CoV-2 Infection Impairs Dendritic Cell and T Cell Responses. Immunity 53, 864–877 e865 (2020).

7. N. Le Bert et al., SARS-CoV-2-specific T cell immunity in cases of COVID-19 and SARS, and uninfected controls. Nature 584, 457–462 (2020).

8. A. Nelde et al., SARS-CoV-2-derived peptides define heterologous and COVID-19-induced T cell recognition. Nat Immunol 22, 74–85 (2021).

9. A. E. Oja et al., Divergent SARS-CoV-2-specific T-and B-cell responses in severe but not mild COVID-19 patients. Eur J Immunol 50, 1998–2012 (2020).

10. Y. Peng et al., Broad and strong memory CD4(+) and CD8(+) T cells induced by SARS-CoV-2 in UK convalescent individuals following COVID-19. Nat Immunol 21, 1336–1345 (2020).

11. A. Grifoni et al., Targets of T Cell Responses to SARS-CoV-2 Coronavirus in Humans with COVID-19 Disease and Unexposed Individuals. Cell 181, 1489–1501 e1415 (2020).

12. T. Sekine et al., Robust T Cell Immunity in Convalescent Individuals with Asymptomatic or Mild COVID-19. Cell 183, 158–168 e114 (2020).

13. H. Kared et al., CD8+ T cell responses in convalescent COVID-19 individuals target epitopes from the entire SARS-CoV-2 proteome and show kinetics of early differentiation. bioRxiv, (2020).

14. A. P. Ferretti et al., Unbiased Screens Show CD8(+) T Cells of COVID-19 Patients Recognize Shared Epitopes in SARS-CoV-2 that Largely Reside outside the Spike Protein. Immunity 53, 1095–1107 e1093 (2020).

15. L. Ni et al., Detection of SARS-CoV-2-Specific Humoral and Cellular Immunity in COVID-19 Convalescent Individuals. Immunity 52, 971–977 e973 (2020).

16. J. Braun et al., SARS-CoV-2-reactive T cells in healthy donors and patients with COVID-19. Nature 587, 270–274 (2020).

17. J. Mateus et al., Selective and cross-reactive SARS-CoV-2 T cell epitopes in unexposed humans. Science 370, 89–94 (2020).

18. S. G. Reed, Cross-viral protection against SARS-CoV-2? Nat Rev Immunol 21, 3 (2021).

19. D. K. Meyerholz, S. Perlman, Does common cold coronavirus infection protect against severe SARS-CoV-2 disease? J Clin Invest 131, (2021).

20. K. Balz, L. Trassl, V. Hartel, P. P. Nelson, C. Skevaki, Virus-Induced T Cell-Mediated Heterologous Immunity and Vaccine Development. Front Immunol 11, 513 (2020).

21. I. J. Amanna, M. K. Slifka, Successful Vaccines. Curr Top Microbiol Immunol 428, 1–30 (2020).

22. S. J. Sung, Monocyte-Derived Dendritic Cells as Antigen-Presenting Cells in T-Cell Proliferation and Cytokine Production. Methods Mol Biol 2020, 131–141 (2019).

23. T. L. Tang-Huau, E. Segura, Human in vivo-differentiated monocyte-derived dendritic cells. Semin Cell Dev Biol 86, 44–49 (2019).

24. C. V. Harding, D. Canaday, L. Ramachandra, Choosing and preparing antigen-presenting cells. Curr Protoc Immunol Chapter 16, Unit 16 11 (2010).

25. V. Mysore et al., FcgR engagement reprograms neutrophils into antigen cross-presenting cells that elicit acquired immunity. Nat Commun In revision, (2021).

26. B. P. Nicolet, A. Guislain, M.C. Wolkers, Combined Single-Cell Measurement of Cytokine mRNA and Protein Identifies T Cells with Persistent Effector Function. J Immunol 198, 962–970 (2017).

27. E. Cano-Gamez et al., Single-cell transcriptomics identifies an effectorness gradient shaping the response of CD4(+) T cells to cytokines. Nat Commun 11, 1801 (2020).

28. I. Thevarajan et al., Breadth of concomitant immune responses prior to patient recovery: a case report of non-severe COVID-19. Nat Med 26, 453–455 (2020).

29. C. Huang et al., Clinical features of patients infected with 2019 novel coronavirus in Wuhan, China. Lancet 395, 497–506 (2020).

30. Z. He et al., Seroprevalence and humoral immune durability of anti-SARS-CoV-2 antibodies in Wuhan, China: a longitudinal, population-level, cross-sectional study. Lancet 397, 1075–1084 (2021).

31. M. Norman et al., Ultrasensitive high-resolution profiling of early seroconversion in patients with COVID-19. Nat Biomed Eng 4, 1180–1187 (2020).

32. J. M. Curtsinger, M. F. Mescher, Inflammatory cytokines as a third signal for T cell activation. Curr Opin Immunol 22, 333–340 (2010).

33. P. Deshpande et al., IL-7-and IL-15-mediated TCR sensitization enables T cell responses to self-antigens. J Immunol 190, 1416–1423 (2013).

34. S. Oh, L. P. Perera, D. S. Burke, T. A. Waldmann, J. A. Berzofsky, IL-15/IL-15Ralpha-mediated avidity maturation of memory CD8+ T cells. Proc Natl Acad Sci U S A 101, 15154–15159 (2004).

35. X. Zhou et al., The deubiquitinase Otub1 controls the activation of CD8(+) T cells and NK cells by regulating IL-15-mediated priming. Nat Immunol 20, 879–889 (2019).

36. R. Setoguchi, IL-15 boosts the function and migration of human terminally differentiated CD8+ T cells by inducing a unique gene signature. Int Immunol 28, 293–305 (2016).

37. T. L. Van Belle et al., IL-15 augments TCR-induced CD4+ T cell expansion in vitro by inhibiting the suppressive function of CD25 High CD4+ T cells. PLoS One 7, e45299 (2012).

38. J. Hagen et al., Comparative Multi-Donor Study of IFNgamma Secretion and Expression by Human PBMCs Using ELISPOT Side-by-Side with ELISA and Flow Cytometry Assays. Cells 4, 84–95 (2015).

39. Y. Tian et al., Unique phenotypes and clonal expansions of human CD4 effector memory T cells re-expressing CD45RA. Nat Commun 8, 1473 (2017).

40. Y. Tian, A. Sette, D. Weiskopf, Cytotoxic CD4 T Cells: Differentiation, Function, and Application to Dengue Virus Infection. Front Immunol 7, 531 (2016).

41. N. Graham et al., Rapid Induction and Maintenance of Virus-Specific CD8(+) TEMRA and CD4(+) TEM Cells Following Protective Vaccination Against Dengue Virus Challenge in Humans. Front Immunol 11, 479 (2020).

42. Y. Tian et al., Dengue-specific CD8+ T cell subsets display specialized transcriptomic and TCR profiles. J Clin Invest 129, 1727–1741 (2019).

43. J. W. Northfield et al., Human immunodeficiency virus type 1 (HIV-1)-specific CD8+ T(EMRA) cells in early infection are linked to control of HIV-1 viremia and predict the subsequent viral load set point. J Virol 81, 5759–5765 (2007).

44. J. M. Dan et al., Immunological memory to SARS-CoV-2 assessed for up to 8 months after infection. Science 371, (2021).

45. A. D. Amir el et al., viSNE enables visualization of high dimensional single-cell data and reveals phenotypic heterogeneity of leukemia. Nat Biotechnol 31, 545–552 (2013).

46. J. G. Burel et al., An Integrated Workflow To Assess Technical and Biological Variability of Cell Population Frequencies in Human Peripheral Blood by Flow Cytometry. J Immunol 198, 1748–1758 (2017).

47. P. A. Szabo et al., Single-cell transcriptomics of human T cells reveals tissue and activation signatures in health and disease. Nat Commun 10, 4706 (2019).

48. Q. Han et al., Polyfunctional responses by human T cells result from sequential release of cytokines. Proc Natl Acad Sci U S A 109, 1607–1612 (2012).

49. B. P. Nicolet et al., CD29 identifies IFN-gamma-producing human CD8(+) T cells with an increased cytotoxic potential. Proc Natl Acad Sci U S A 117, 6686–6696 (2020).

50. V. S. Patil et al., Precursors of human CD4(+) cytotoxic T lymphocytes identified by single-cell transcriptome analysis. Sci Immunol 3, (2018).

51. A. Gil et al., Vaccination and heterologous immunity: educating the immune system. Trans R Soc Trop Med Hyg 109, 62–69 (2015).

52. I. Lopez-Martin, E. Andres Esteban, F.J. Garcia-Martinez, Relationship between MMR vaccination and severity of Covid-19 infection. Survey among primary care physicians. Med Clin (Engl Ed) 156, 140–141 (2021).

53. J. E. Gold et al., Analysis of Measles-Mumps-Rubella (MMR) Titers of Recovered COVID-19 Patients. mBio 11, (2020).

54. F. Lievano et al., Measles, mumps, and rubella virus vaccine (M-M-RII): a review of 32 years of clinical and postmarketing experience. Vaccine 30, 6918–6926 (2012).

55. A. Mantovani, M. G. Netea, Trained Innate Immunity, Epigenetics, and Covid-19. N Engl J Med 383, 1078–1080 (2020).

56. B. Agrawal, Heterologous Immunity: Role in Natural and Vaccine-Induced Resistance to Infections. Front Immunol 10, 2631 (2019).

57. L. A. J. O’Neill, M. G. Netea, BCG-induced trained immunity: can it offer protection against COVID-19? Nat Rev Immunol 20, 335–337 (2020).

58. C. Covian et al., BCG-Induced Cross-Protection and Development of Trained Immunity: Implication for Vaccine Design. Front Immunol 10, 2806 (2019).

59. M. G. Netea et al., Trained Immunity: a Tool for Reducing Susceptibility to and the Severity of SARS-CoV-2 Infection. Cell 181, 969–977 (2020).

60. L. F. Su, B. A. Kidd, A. Han, J. J. Kotzin, M. M. Davis, Virus-specific CD4(+) memory-phenotype T cells are abundant in unexposed adults. Immunity 38, 373–383 (2013).

61. J. C. Nolz, M. J. Richer, Control of memory CD8(+) T cell longevity and effector functions by IL-15. Mol Immunol 117, 180–188 (2020).

62. T. P. Arstila et al., A direct estimate of the human alphabeta T cell receptor diversity. Science 286, 958–961 (1999).

63. L. Wooldridge et al., A single autoimmune T cell receptor recognizes more than a million different peptides. J Biol Chem 287, 1168–1177 (2012).

64. C. H. Coles et al., TCRs with Distinct Specificity Profiles Use Different Binding Modes to Engage an Identical Peptide-HLA Complex. J Immunol 204, 1943–1953 (2020).

65. S. Sorup et al., Simultaneous vaccination with MMR and DTaP-IPV-Hib and rate of hospital admissions with any infections: A nationwide register based cohort study. Vaccine 34, 6172–6180 (2016).

66. B. Beric-Stojsic, J. Kalabalik-Hoganson, D. Rizzolo, S. Roy, Childhood Immunization and COVID-19: An Early Narrative Review. Front Public Health 8, 587007 (2020).

67. C. Pawlowski et al., Exploratory analysis of immunization records highlights decreased SARS-CoV-2 rates in individuals with recent non-COVID-19 vaccinations. Sci Rep 11, 4741 (2021).

68. Q. X. Long et al., Immune memory in convalescent patients with asymptomatic or mild COVID-19. Cell Discov 7, 18 (2021).

69. L. E. Thomas, F. Li, M.J. Pencina, Overlap Weighting: A Propensity Score Method That Mimics Attributes of a Randomized Clinical Trial. JAMA 323, 2417–2418 (2020).

70. G. X. Zheng et al., Massively parallel digital transcriptional profiling of single cells. Nat Commun 8, 14049 (2017).

71. T. Stuart et al., Comprehensive Integration of Single-Cell Data. Cell 177, 1888–1902 e1821 (2019).

72. L. Jehi et al., Individualizing Risk Prediction for Positive Coronavirus Disease 2019 Testing: Results From 11,672 Patients. Chest 158, 1364–1375 (2020).

73. N. Mehta et al., Association of Use of Angiotensin-Converting Enzyme Inhibitors and Angiotensin II Receptor Blockers With Testing Positive for Coronavirus Disease 2019 (COVID-19). JAMA Cardiol 5, 1020–1026 (2020).

